# Group I p21-activated kinases in leukemia cell adhesion to fibronectin

**DOI:** 10.1101/2020.08.25.266056

**Authors:** Kateřina Kuželová, Adam Obr, Pavla Röselová, Dana Grebeňová, Petra Otevřelová, Barbora Brodská, Aleš Holoubek

## Abstract

P21-activated kinases (PAK) regulate many processes associated with cytoskeleton dynamics, including cell adhesion, migration, and apoptosis. PAK function is frequently altered in cancer, and PAK were proposed as therapy targets both in solid tumors and in hematological malignancies. However, current knowledge about PAK function in cell adhesion is mainly based on adherent cell models. Moreover, existing functional differences among the individual PAK family members are unsufficiently characterized.

We measured expression of PAK group I members in leukemia cell lines and in primary leukemia cells, both on protein and mRNA levels. In functional assays, we analyzed the effect of two PAK inhibitors with different mechanisms of action, IPA-3 and FRAX597. Changes in cell interaction with fibronectin were monitored through impedance measurement and by interference reflection microscopy. Cytotoxic effects of inhibitors were assessed by Annexin V/propidium iodide test. PAK intracellular localization was analyzed by confocal microscopy.

PAK2 transcript was dominant in cell lines, whereas primary leukemia cells also expressed comparable amount of PAK1, which was detected as two transcription isoforms: PAK1-full and PAK1Δ15. PAK1Δ15 and PAK2 transcript levels correlated with surface density of integrins β1 and αVβ3. PAK1-full, but not PAK2, was present in membrane protrusions. The inhibitors had partly opposed effects: IPA-3, which prevents PAK activation, induced cell contraction in semi-adherent HEL cells only. FRAX597, which inhibits PAK kinase activity, increased cell-surface contact area in all leukemia cells. Both inhibitors reduced the stability of cell attachment and induced cell death. Although many cells accumulated high FRAX597 amounts, low doses were sufficient to kill sensitive cells. FRAX597-induced cell death was fast in the MV4-11 cell line and in primary AML cells.

Although PAK group I seem to be essential for leukemia cell adhesion and survival, and might thus serve as therapy targets, many PAK functions still remain to be attributed to individual isoforms and to their functional domains.

## Background

Binding of hematopoietic stem and progenitor cells to the extracellular matrix of the bone marrow (BM) provides important intracellular signals, which regulate multiple processes, including self-renewal, proliferation, and differentiation of these cells. In leukemia, interaction with the BM microenvironment promotes resistance to chemotherapy and survival of malignant cells (1, 2). Acute myeloid leukemia (AML) is still associated with high mortality, which is due in particular to relapses from chemotherapy-resistant leukemia stem cells. Interference with adhesion signaling could release these cells from their niches and enhance the efficiency of chemotherapy. However, in contrast to adherent cell types, adhesion signaling in stem hematopoietic/leukemia cells is not well described. Cell-matrix contact points are formed by large protein complexes similar to those of immature focal adhesions occurring in adherent cell types (3). Leukemia cell adhesion structures lack typical markers of mature focal adhesions, such as talin, paxillin, and vinculin. They are not connected to actin fibers, and are in general more dynamic than focal adhesions in adherent cells.

P21-activated kinases (PAK) regulate diverse processes involving cytoskeletal rearrangements, such as cell adhesion and migration, cell division, or cell-cell interactions. They are considered as therapeutic targets in different cancer types including hematological malignancies (4-9). Possible role of PAK in tumorigenesis has been reviewed (10) with a focus on PAK1 and PAK4.

In adherent cells, the switch between adherent and migratory phenotype largely depends on the activity of c-Src, and at least one of the six known PAK family members, PAK1, is an important c-Src downstream effector (11-14). However, the role of other members of PAK family in mediating c-Src regulatory effects in adherent cells is not clear. PAK1 belongs to a group of proteins with serine/threonine kinase activity as well as with non-kinase functions, and PAK group I includes three proteins (PAK1 to PAK3) with high sequence homology. At least one of them, PAK2, directly interacts with c-Src according to Human Protein Reference Database (15). Phosphorylation by Src was also shown to prime PAK2 for activation by small GTPases (16).

We have found previously that the stability of adhesion structures in leukemia cells is not significantly affected by c-Src inhibition (17), probably due to the lack of their connection to actin fibers. On the other hand, we and others have shown that PAKs are required for leukemia cell binding to fibronectin, and also for their survival (5, 18). PAK3 is only found in a few cell types, but PAK2 is ubiquitously expressed, and it was reported to have important roles in hematopoietic cells (19-22). PAK group I are also involved in leukemogenesis driven by aberrantly activated protein kinases (23-25). Despite virtually identical substrate specificity of PAK1 and PAK2 kinase domains (26), these proteins have at least partly non-redundant functions (27, 28). In addition, PAK1 has many transcription isoforms, and at least two of them are translated (NCBI Reference Sequence NM_001128620.2 and NM_001376273.1). We have recently described their protein products, the full-length PAK1 (PAK1-full) and a shorter isoform, PAK1Δ15, formed from alternatively spliced mRNA lacking the exon 15. In adherent cells, PAK2 and PAK1Δ15 were enriched in adhesion sites, whereas PAK1-full was required for cell spreading (29). Moreover, all these PAK isoforms form homo- and heterodimers and likely regulate each other (29). Given the importance of cell-matrix interactions for leukemia persistence, we aimed to analyze in more detail the role of PAK in leukemia cell adhesion. In this work, we analyzed PAK group I expression in leukemia cell lines and in primary cells of AML patients, and we correlated these data with cell adhesivity to fibronectin as well as with expression of selected integrins. To determine the type of PAK activity (kinase or non-kinase), which is required for cell-matrix interaction and for cell survival, we compared the effects of two PAK inhibitors with different mechanisms of action, IPA-3 and FRAX597. Changes in cell binding to fibronectin-coated surface were monitored in real-time using electric cell-substrate impedance sensing (ECIS). This method allowed us to detect differences in the acute cell response to IPA-3 versus FRAX597. The cell-surface contact area was visualized by interference reflection microscopy (IRM). We combined these analyses with measurement of adherent cell fraction (ACF), which is indicative of the stability of cell attachment.

## Methods

### Cell isolation and culture

OCI-AML3, OCI-AML2, MV4-11, KG-1, and Kasumi-1 cell lines were purchased from DSMZ (Germany), HL-60 cells from ECACC (Salisbury, UK). After receipt, the cells were shortly expanded and aliquots were cryopreserved for later use. Freshly thawed aliquots were used for experiments without authentication. HEL cells were obtained as a gift and authenticated using analysis of short tandem repeats, the results were compared with ATCC database. The cell line MOLM-7 (not commercially available) was obtained from the laboratory of origin (30). The majority of cell lines were cultured in RPMI-1640 medium with 10 % fetal calf serum (FCS), 100 U/ml penicillin, and 100 µg/ml streptomycin at 37 °C in 5 % CO_2_ humidified atmosphere, except for OCI-AML2 and OCI-AML3 (alpha-MEM medium, 20 % FCS), and Kasumi-1 (RPMI medium, 20 % FCS).

Primary cells from AML patients were obtained from leukapheresis at diagnosis. All patients provided their written informed consent as to the use of their biological material for research purposes, in accordance with the Helsinki Declaration. The project was approved by the Ethics Commitee of the Institute of Hematology, and all experiments were performed in accordance with relevant guidelines and regulations. The leukapheretic products were diluted 20-fold in phosphate buffered saline (PBS) and the mononuclear cell fraction was then separated using Histopaque-1077 (Sigma, #H8889). Analysis by flow cytometry (CD45/SSC dotplots) confirmed a high prevalence of leukemia blasts in mononuclear cell samples.

### Chemicals and antibodies

Fibronectin fragment (120 kDa cell attachment region) was from Merck (F1904). IPA-3 (#3622) and PIR3.5 (#4212) were purchased from Tocris Bioscience and dissolved in sterile dimethylsulfoxide (DMSO) to make 50 mM stock solution. Working solutions were prepared by 10-fold dilution of the stock solution in 50 mM Tris, pH 8.0, immediately before use. FRAX597 (S7271) and dasatinib (S1021) were obtained from SelleckChem, stock solutions were made in DMSO at 10 mM or 200 µM concentration, respectively, and diluted in culture media immediately before use. In all experiments, the cell density was adjusted to 3×10^5^ cells/ml prior to treatment with inhibitors.

The antibodies used were the following: PAK1 (Cell Signaling, #2602), PAK1 (Abcam, ab223849), PAK2 (Abcam, #ab76293), PAK1 (phosphoS144)+PAK2 (phosphoS141)+PAK3 (phosphoS139) (Abcam, #ab40795), β-actin (Sigma-Aldrich, A5441), CD45-V450 (BD Biosciences, #560367), CD34-BV786 (BD Biosciences, #743534), Integrin β1-AlexaFluor647 (Abcam, ab193592), Integrin αVβ3-FITC (Abcam, ab93513). All PAK antibodies were previously characterized using siRNA (29).

### Real-Time Polymerase Chain Reaction

RNA from 2×10^7^ cells was isolated using the RNeasy Mini Kit (Qiagen, Venlo, Netherlands), cDNA was generated by reverse transcription on CFX96 real-time system (BioRad) using SensiFAST cDNA Synthesis Kit (Bioline, London, UK). Template RNA and resulting cDNA quality and concentration were assessed using ND-1000 Nanodrop system (Thermo Fisher Scientific, Waltham, MA, USA). The relative amount of PAK transcript variants was measured by quantitative real-time PCR (qPCR) using SensiFAST SYBR No-ROX Kit (Bioline) and calculated using Bio-Rad CFX Manager Software. Primers were designed to target the full-length PAK1 (transcript variant 1, NCBI RefSeq NM_001128620.2), PAK1Δ15 (transcript variant 8, NCBI RefSeq NM_001376273.1), and PAK2 (NM_002577.4). The sequences of the primer pairs were the following: PAK1 full-length: forward GCTACAGGTGAGAAAACTGAGG, reverse CTCACAGAAGGCTTGGCACA, PAK1Δ15: forward GAGCTGCTACAGCATCAATTCC, reverse GCAAGGAGAAGAGGGCATCA, PAK2: forward CCCTTGAGGGCCTTGTACCT, reverse CGGTTTGGCCAGTTTCAGGA. For the relative quantification by 2^−ΔΔCt^ method, GAPDH expression was measured as a reference, using GAAACTGTGGCGTGATGGC and CCGTTCAGCTCAGGGATGAC as the forward and reverse primer, respectively. The primers were designed to work at identical conditions, which allowed us to perform PCR reactions for all PAK variants simultaneously.

### Western-blotting

The cells were pelleted by centrifugation and lysed for 30 min/4 °C in modified RIPA lysis buffer (50 mM HEPES; 0,15 M NaCl; 2 mM EDTA; 0,1 % NP-40; 0,05 % sodium deoxycholate) with freshly added protease and phosphatase inhibitors. Cellular debris was removed by centrifugation (16.000 g/4 °C/15 min), the lysate was mixed 1:1 (v/v) with 2x Laemmli sample buffer and incubated for 5 min at 95 °C. Alternatively, the cell pellets were lysed directly in Laemmli sample buffer, with very similar results.

An equivalent of 20 µg of total protein was resolved on 7.5% polyacrylamide gel (18×18 cm) and transferred to a nitrocellulose membrane. The membrane was incubated with primary antibodies for 1 h at the room temperature, in 3 % BSA in PBS with 0,1 % Tween-20 (PBST), washed in PBST six times, and incubated with corresponding HRP-conjugated secondary antibodies for 1 h. Chemiluminiscence signal was developed by Clarity Western ECL Substrate (BioRad, #170-5060), and detected and analyzed using G:BOX iChemi XT-4 (Syngene).

### Flow-cytometry

To characterize primary mononuclear cell preparations, cells were stained with the appropriate antibodies for 30 min at 5°C, washed once in PBS, and analyzed on a BD Fortessa flow cytometer. The debris were outgated, and blasts and lymphocytes were discriminated in CD45 versus side scatter (SSC) dotplots. CD34 and integrin expression was determined from the cells in the blast gate, as it was described previously (31).

To determine the intracellular amount of FRAX597, we used the fluorescence intensity in the Pacific blue channel (405/450 nm) measured from aliquots of cell line samples without further processing. The mean fluorescence intensity of control (untreated) samples was used as the background and subtracted from the measured values of treated samples.

### Assessment of apoptotic/necrotic cell fraction

Annexin V/propidium iodide staining was performed in leukemia cell lines using a standard protocol after 5h incubation with inhibitors. Primary cell viability was measured by propidium iodide exclusion test from aliquots of cell suspensions without further processing.

### Adherent cell fraction (ACF) measurement

The end-point method for assessment of cellular adhesivity to fibronectin-coated surface has been described previously (32). Briefly, the cells (3×10^4^ in 100 µl) were seeded into fibronectin-coated wells of a microtitration plate and incubated at 37 °C. Then, the suspension was aspirated and the wells were washed with PBS/Ca^2+^/Mg^2+^. The remaining cells were quantified by means of fluorescent labelling (CyQuant Cell Proliferation Assay Kit; Molecular Probes, #C7026). The adherent cell fraction (ACF) was calculated reporting the fluorescence signal from the fibronectin-coated plate to the signal obtained from a reference plate containing the total cell number.

### Electric cell-substrate impedance sensing (ECIS)

Impedance measurements were performed using the ECIS Zθ device (Applied Biophysics). The wells of a 8W10E+ plate were hydrated overnight in culture medium, washed with sterile water, filled with 150 µl FN solution (20 µg/ml in 0.15 M NaCl), incubated for 30 min at the room temperature or overnight at 5 °C, washed with RPMI medium without calf serum and filled with 200 µl medium with serum. After electrode stabilization, the baseline was monitored for at least 1 h before addition of cells (120.000 per well). In the first setting, 200 µl of cell suspension was added to the wells and the cell attachment was monitored for about 1.5 h prior to inhibitor addition. In the second setting, the medium was removed and 400 µl of cell suspension pretreated with inhibitors was added. One of the wells in each plate were left empty (medium only) and the signal was used as the baseline for the other wells. The instrument automatically decomposes the impedance signal into resistance and capacitance. The ECIS records were exported to Excel and processed using the GraphPad Prism software: the background was set to zero at a time point shortly before cell seeding, and the baseline (empty well) was subtracted. The signals shown in the graphs represent the averages from two identically treated wells, which were run in parallel.

### Cell transfection and confocal microscopy

Plasmids coding for fluorescently labeled PAK isoforms were prepared as described previously (29). MOLM-7 cells (6×10^5^) were transfected using the Neon™ transfection system (1325 V, 10 ms, 3 cycles) and 0.5 µg of DNA. The transfected cells were cultured for 24 h in RPMI medium without antibiotics, incubated for 1 h on fibronectin-coated coverslips, and localization of PAK1-full and PAK2 was analyzed by fluorescence microscopy (FV-1000 confocal microscope, Olympus). For analysis of changes in the cell-surface contact area, cells were incubated for 90 min on fibronectin-coated surface, treated for 30 min with 10 µM FRAX597, and analyzed in the interference reflection mode. The measurement was performed by means of FV-1000 confocal microscope (Olympus), using 405 nm laser beam and focusing to the glass surface.

### Statistical analyses

Statistical evaluation of experimental results was performed using the GraphPad Prism 7 software. The p-value limit for statistically significant differences between groups was set to 0.05.

## Results

### PAK expression in leukemia cells

PAK expression was analyzed using previously characterized antibodies against PAK1, PAK2, or the autophosphorylation site pSer144/141 on PAK1/PAK2, respectively (29). The band pattern obtained from leukemia cells was similar to that from adherent cells: PAK2 antibody recognizes a dominant band at 60 kDa, whereas PAK1 is detected in several bands in the range from 64 to 67 kDa (Fig. 1). The bands of the full-length PAK1 (PAK1-full) and of the shorter isoform PAK1Δ15, which were exogenously expressed in HEK-293T cells, are shown in the first two lanes of Fig. 1A. Each PAK1 isoform gives rise to several western-blot bands, probably due to yet unknown posttranslational modification(s). In addition, the apparent molecular weight (MW) of PAK1 is slightly modified by multiple phosphorylation (29). It was thus not possible to attribute the individual endogenous bands to individual PAK1 isoforms. The extent of phosphorylation at pSer144/141 depends on PAK1/PAK2 kinase activity. As we have reported previously (29), the exogenous PAK1 forms are predominantly unphosphorylated. Figure 1B shows PAK expression in AML-derived cell lines in comparison with primary mononuclear cells from AML patients. In general, the amount of phosphorylated PAK1 (pPAK1) compared to pPAK2 was higher in primary cells. MW of PAK1 bands was essentially the same, but the bands at the highest MW were often more intense in primary samples. Fig. 1C documents low PAK1/pPAK1 and high PAK2 levels found in MOLM-7 cell line, which was derived from a patient with chronic myeloid leukemia.

**Figure 1:**
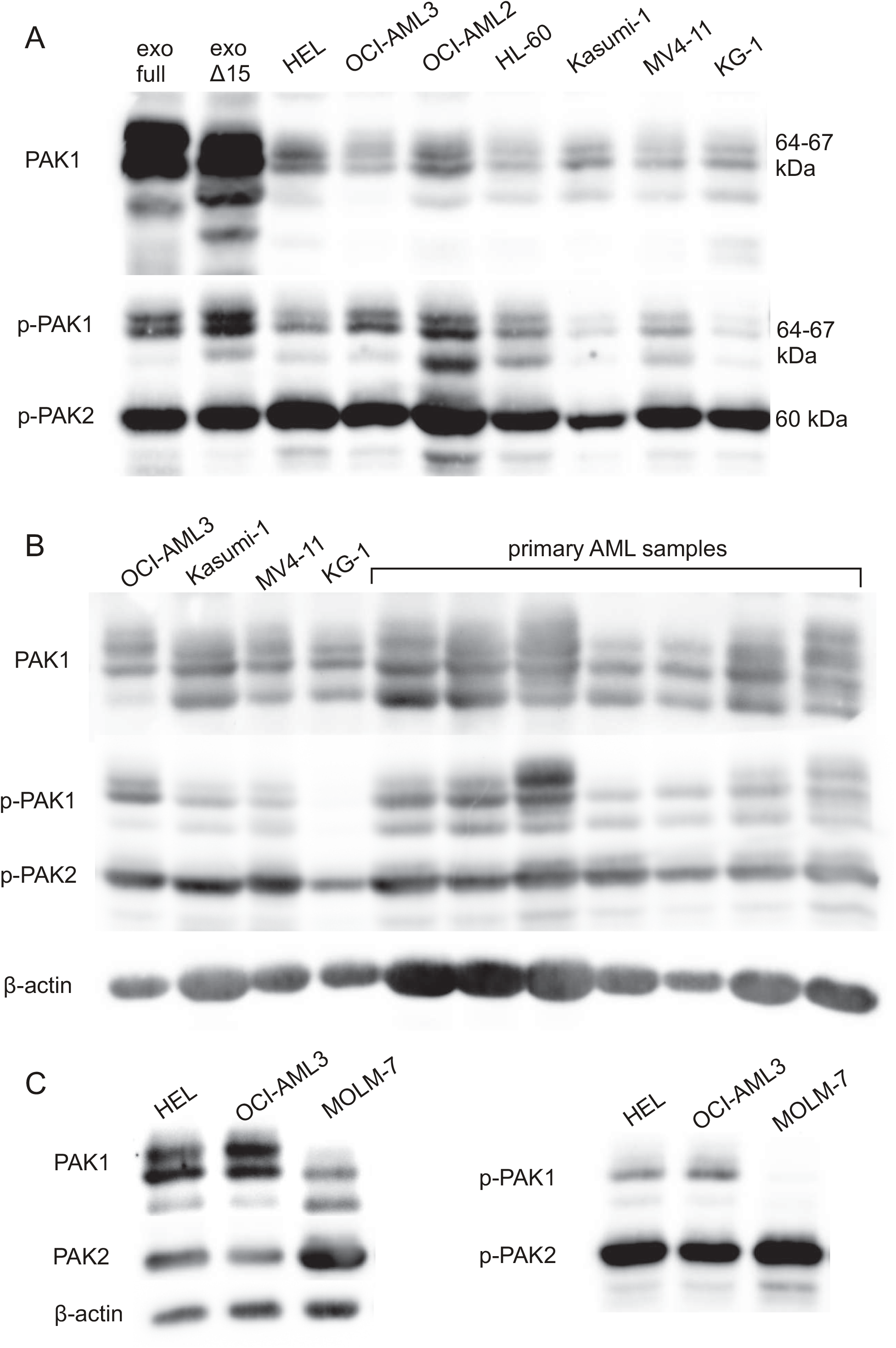
PAK protein expression in leukemia cells. PAK total protein levels and phosphorylation at Ser144/141 of PAK1/PAK2 was assessed in cell lysates from leukemia cell lines, primary AML cells and HEK-293T cells transfected with plasmids for exogenous expression of the full-length PAK1 or PAK1Δ15. A: The exogenously expressed PAK1 is shown in the first two lanes for comparison of the specific PAK1-full and PAK1Δ15 bands with the endogenous bands from leukemia cell lines. B: Comparison of PAK bands in AML cell lines (first four lanes) with those in primary AML samples. C: PAK expression in the cell lines, that were further studied in detail. β-actin was used as a loading control.

PAK1-full and PAK1Δ15 can be distinguished at the transcript level and we performed the analysis both in leukemia cell lines and in primary cells. The PrimerBLAST tool predicts additional PAK1 transcript variants to be detected by the designed primer pairs for PAK1-full and PAK1Δ15. All the five alternative transcript variants for PAK1-full encode the same protein product with 553 aminoacids. Also, two-thirds (16 of 24) of alternative transcripts for PAK1Δ15 share the coding sequence with the reference variant NM_001376273.1, which encodes a protein consisting of 545 aminoacids. The remaining protein variants, which could be produced from 8 alternative PAK1Δ15 transcripts, differ in their N-terminal parts, whereas the C-terminal part, which constitutes the difference between PAK1-full and PAK1Δ15, is conserved. The primers thus correctly reflect the relative abundance of PAK1 variants lacking the exon 15, and thus resembling to PAK2 at their C-terminal part (33). PAK2 primers should only detect the full-length PAK2 transcript, and no cross-recognition was predicted between PAK1 and PAK2 primers. Both PAK1-full and PAK1Δ15 transcripts were more abundant in primary cells (Fig. 2B) compared to cell lines (Fig. 2A), whereas PAK2 levels were similar. PAK1Δ15 variants were expressed at significantly higher levels than PAK1-full (Fig. 2C).

**Figure 2:**
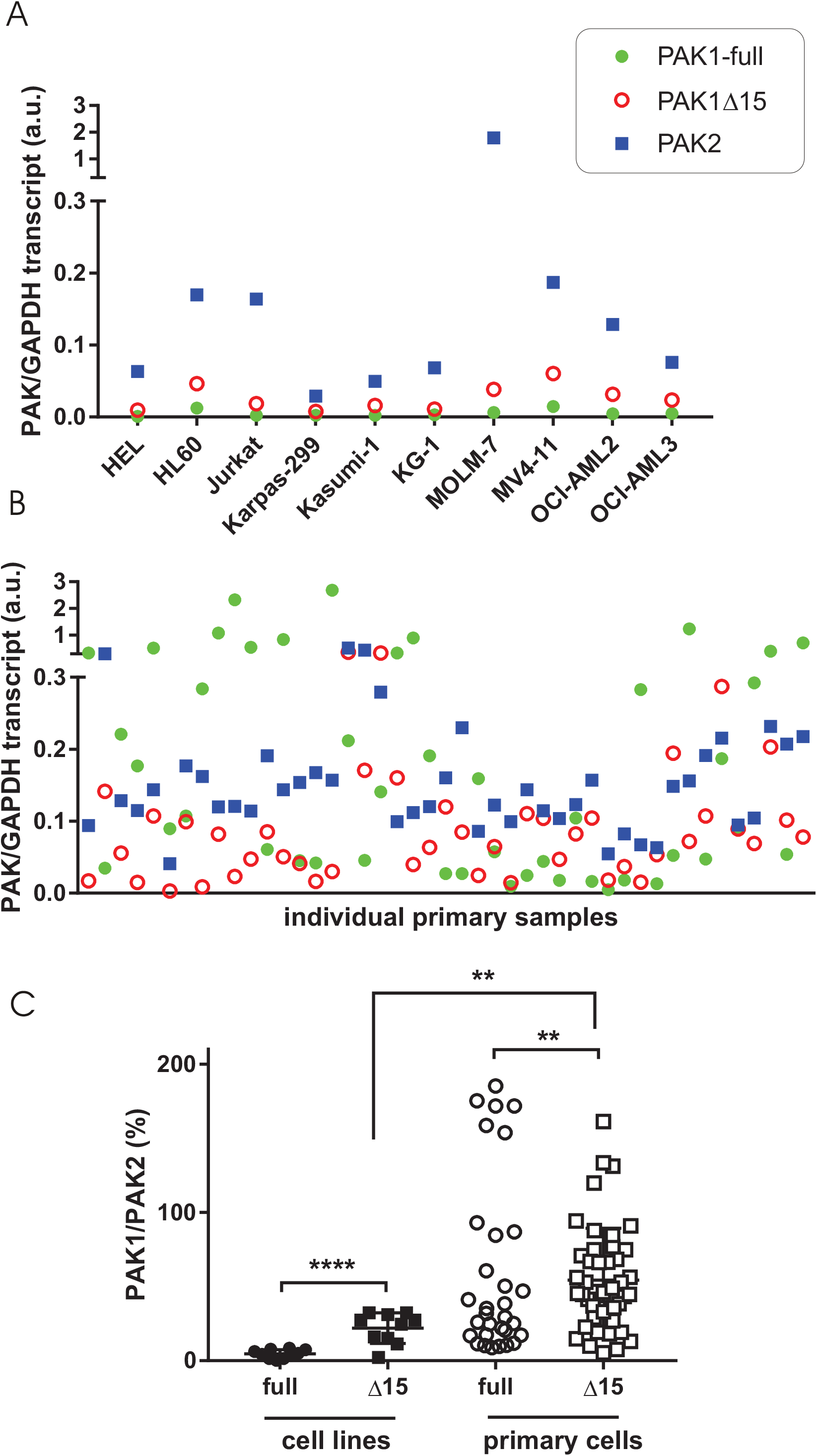
Transcript levels for PAK1-full, PAK1Δ15, and PAK2 in leukemia cell lines and in primary cells. A-B: Values for the individual PAK isoforms normalized to GAPDH in cell lines (A) and in primary cells from 45 AML patients (B). Green closed circles: PAK1-full, red open circles: PAK1Δ15, blue squares: PAK2. C: Summary results for PAK1-full and PAK1Δ15, expressed as relative to PAK2 levels. Differences between the two PAK1 isoforms and between cell lines and primary cells were assessed using standard two-tailed t-test: ** p less than 0.01, **** p less than 0.0001.

### PAK intracellular localization

We have previously uncovered differences in intracellular localization of PAK group I isoforms in adherent cell lines (29). PAK1Δ15 and PAK2 were enriched in adhesion structures, whereas PAK1-full was found in membrane protrusions at the cell periphery. The work with leukemia cells was more challenging due to a very low fraction of cells suitable for microscopic analysis. Cell transfection with plasmids encoding fluorescently labeled PAK largely altered cell viability and adhesivity to fibronectin, although no problem occurred with a control GFP-only plasmid. Nevertheless, in cells with normal morphology and with both green and red fluorescence signals, the images were in line with our previous findings. Leukemia cells do not form mature focal adhesions and the distribution of PAK1-full and PAK2 was rather uniform. However, the signal from PAK1-full was clearly present in thin membrane protrusions, which did not contain PAK2 (Fig. 3).

**Figure 3:**
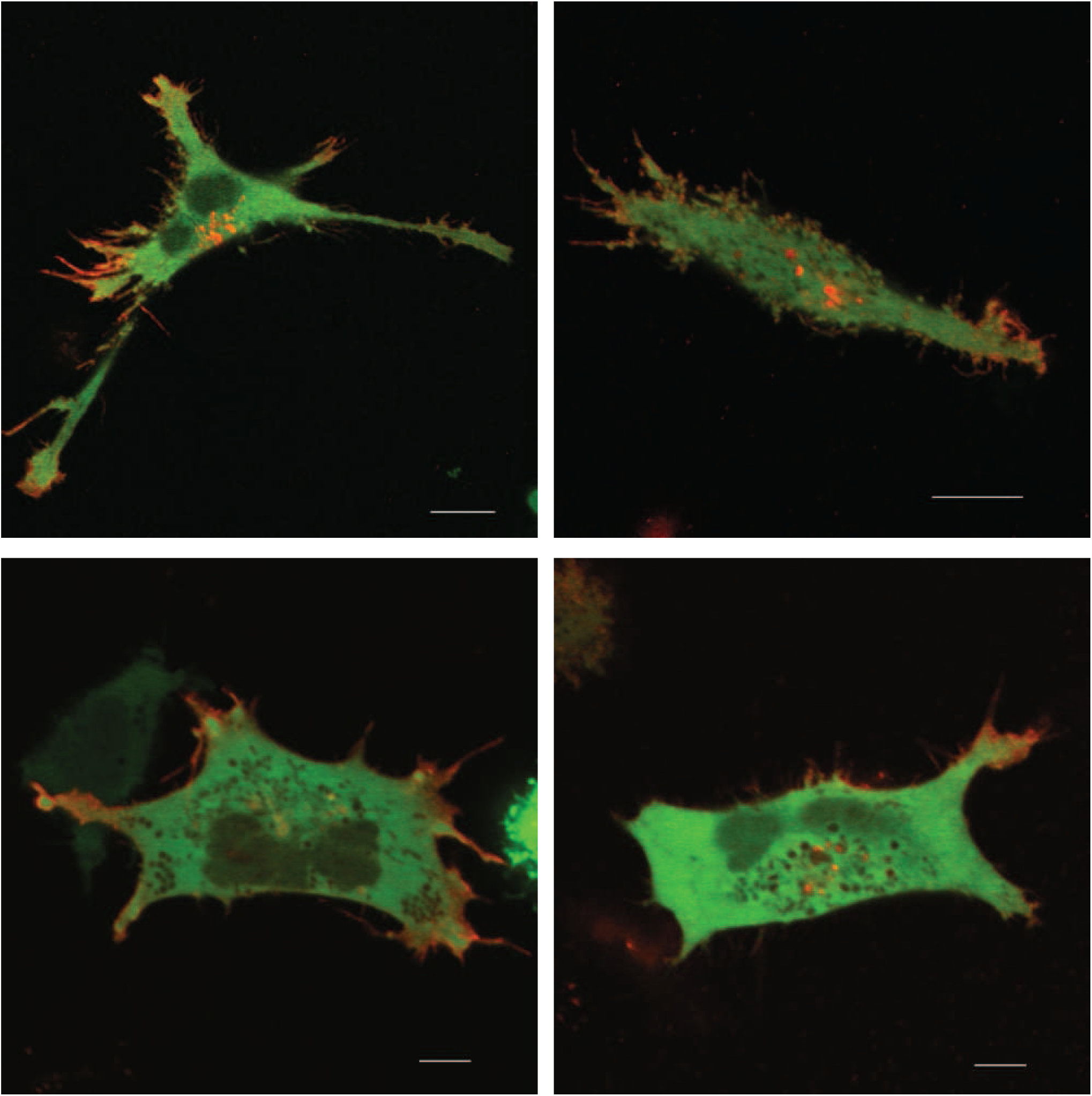
Intracellular localization of PAK1-full and PAK2 in leukemia cells. MOLM-7 cells were co-transfected with plasmids coding for fluorescently labeled PAK1-full (mCherry, red) and PAK2 (eGFP, green), seeded on fibronectin, and the subcellular distribution of the fluorescent proteins was analyzed by confocal microscopy. Scale bars: 10 µM.

### Inhibitor characterization

Previous reports indicated that PAK inhibition by a specific compound, IPA-3, was associated with apoptosis induction (5) as well as with loss of adhesivity to fibronectin in leukemia cell lines (34). We compared the effect of IPA-3 to that of FRAX597, which has different mechanism of action: IPA-3 binds to PAK in their inactive conformation and prevents activation by allosteric changes (35), whereas FRAX597 competes with ATP and inhibits PAK kinase activity (36), presumably without affecting non-kinase functions. PIR3.5 is a structural isomer of IPA-3 with no inhibitory activity against PAK (35), and it is usually used as a negative control compound to check for possible unspecific effects, such as reactive oxygen species generation by IPA-3. The efficiency of IPA-3 depends on the cell density, and we thus adhered to identical conditions in all experiments: the cell suspension was diluted to 3×10^5^ cells/ml at the time point of IPA-3 treatment. Dasatinib was included as a potent inhibitor of kinases of the Src family, which are known to regulate some of PAK functions, at least in adherent cells. It was used at a high (100 nM) concentration inducing a rapid and complete inhibition of Src kinase activity (17).

The efficiency of IPA-3 and FRAX597 in inhibiting PAK kinase activity was compared using the extent of dephosphorylation at Ser144/141 after 1h cell treatment (examples are given in Supplementary Figure S1). In agreement with our previous results in adherent cells (29), one of the phospho-PAK1 bands, denoted as pPAK1-low, was reduced at lower IPA-3 doses then the others. The intensity of pPAK1-low in the cell lines decreased by at least 90% after 1h treatment with 20 µM IPA-3, whereas the other bands were much less reduced (Fig. 4, left). In MOLM-7 cells, only PAK2 phosphorylation was evaluated as the intensity of pPAK1 bands was very low (Fig. 1C). IPA-3 treatment was associated with extensive cell death in many cell lines (Fig. 5) and we thus set the maximal IPA-3 concentration to 50 µM. As we expected, FRAX597 was more efficient in reducing Ser144/141 phosphorylation and its effect was similar for all PAK forms (Fig. 4, right). EC50 was 2 µM or less for all the studied cell types, including primary cells. For subsequent experiments, we used IPA-3 and PIR3.5 at 20 and 50 µM, and FRAX597 at 2 and 10 µM.

**Figure 4:**
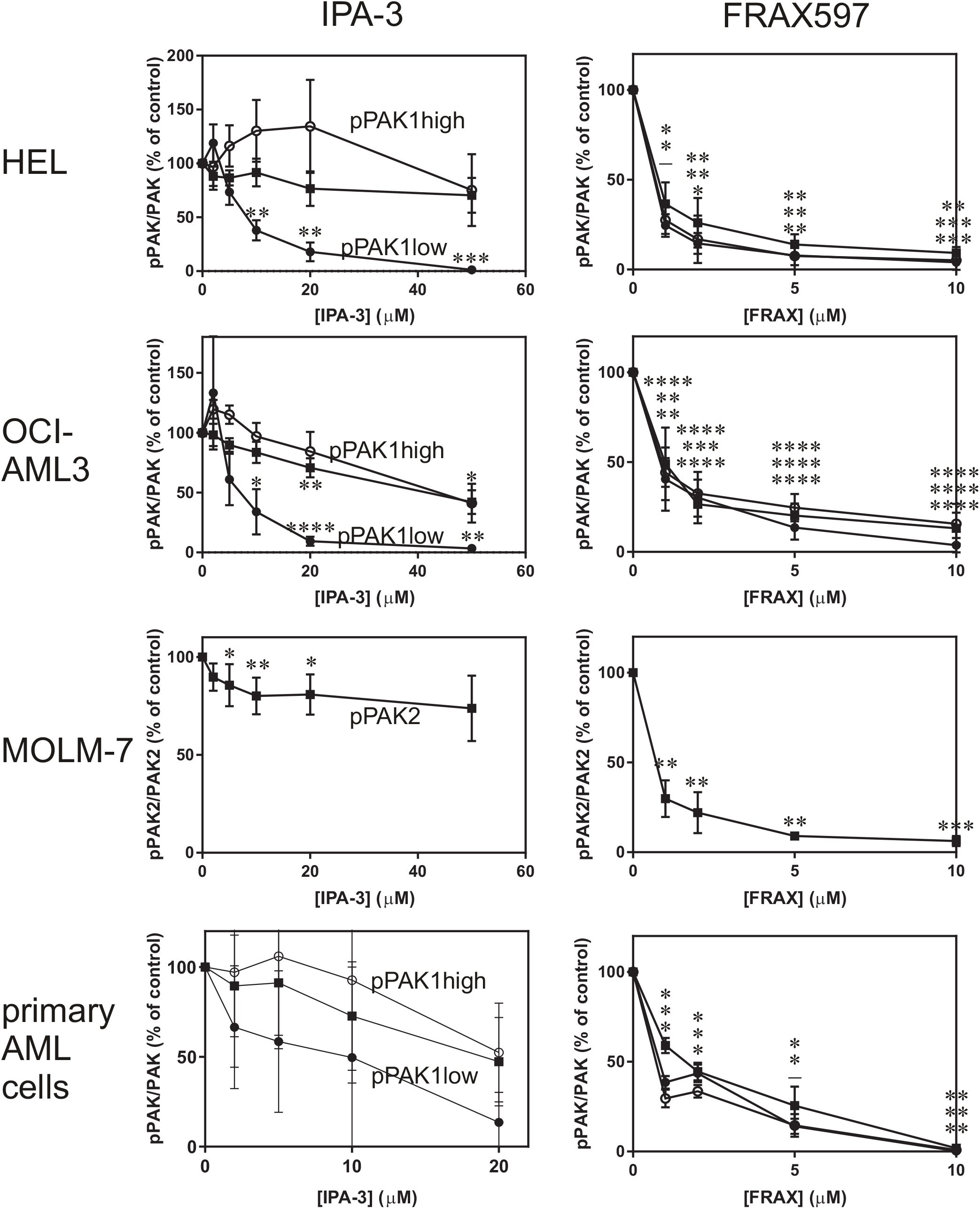
Effect of PAK inhibitors on PAK1/PAK2 phosphorylation at Ser144/141. Cells were treated for 1h with IPA-3 (left) or FRAX597 (right) at different concentrations as indicated, and the amount of pSer144/141 and of total PAK1 and PAK2 was determined by western-blotting. Summary results from at least three experiments are shown for pPAK1 high (open circles), pPAK1 low (closed circles) and pPAK2 (closed squares) normalized to the corresponding total PAK. In MOLM-7 cells, only pPAK2 was evaluated due to very low amount of pPAK1. Individual examples of western-blots are given in the Supplementary Fig. S1. The last row shows summary results from different primary cell samples. Paired t-test was used to assess statistical significance of the difference between treated samples and the corresponding controls (* p < 0.05, ** p < 0.01, *** p < 0.001, **** p < 0.0001). In the right column (FRAX597), the significance indicators are given always in the same order for pPAK1high, pPAK1low, and pPAK2.

**Figure 5:**
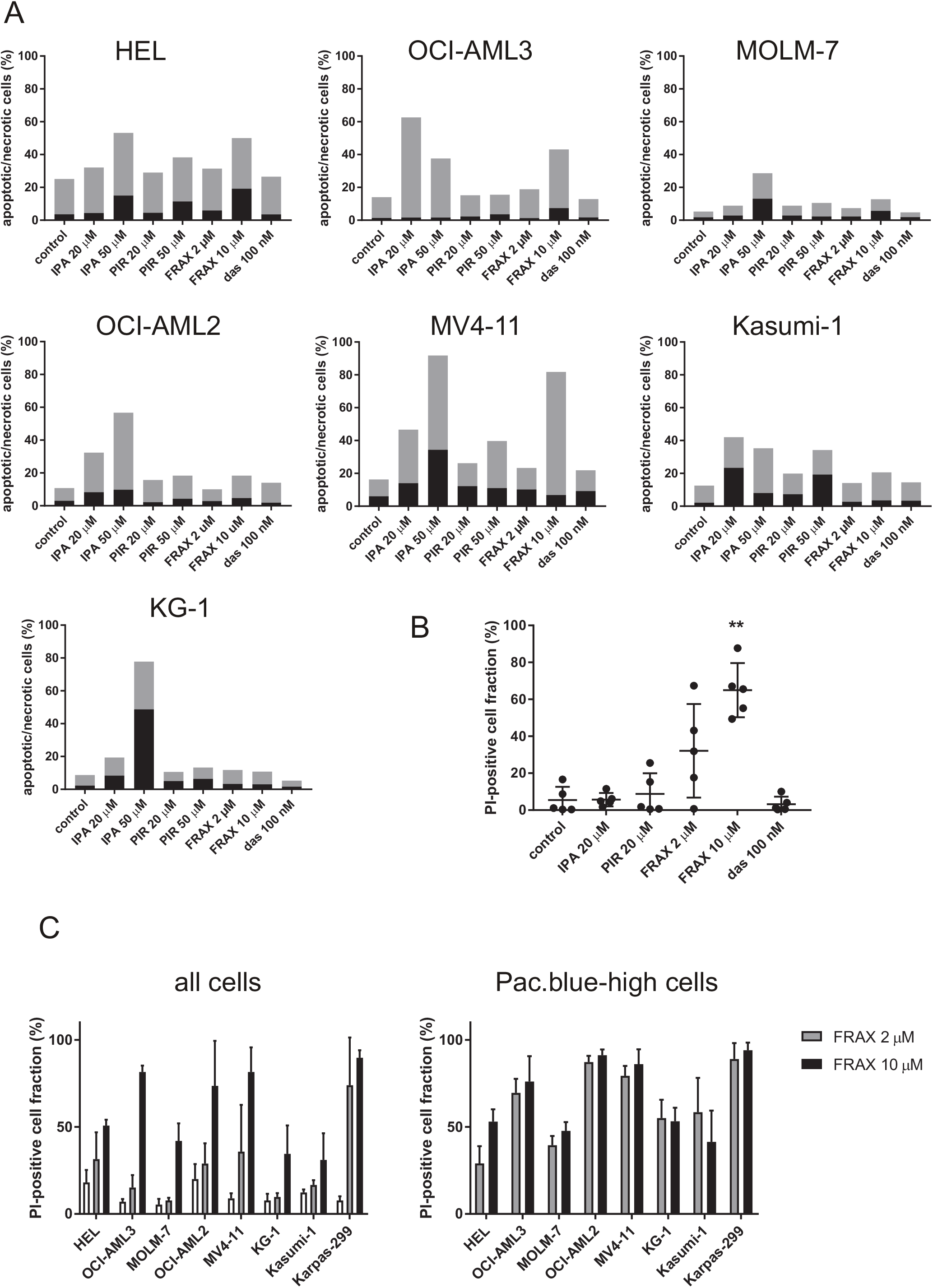
Toxicity of inhibitors in leukemia cells. A: Cell line samples were treated with IPA-3, PIR3.5, FRAX597, or dasatinib for 5 h, and the amount of apoptotic/necrotic cells was assessed by Annexin V/propidium iodide staining. Black parts: early apoptotic cells (Annexin V-positive, PI-negative), grey parts: late apoptotic/necrotic cells (PI-positive). Representative examples from at least two independent experiments for each cell line. B: Viability of primary AML cells after 5h treatment with inhibitors was assessed by propidium iodide staining. The graph shows individual values from experiments with cells from five different AML patients, the means and the standard deviations. The difference between controls and samples treated with 10 µM FRAX597 was statistically significant (p less than 0.01 from standard paired two-tailed t-test). C: Cells were treated for 22 h with 2 or 10 µM FRAX597, and their viability was assessed by propidium iodide staining. Left: viability in the whole cell population, right: viability in cells with high FRAX597 content (gated in the Pacific blue channel). White bars: controls, gray bars: 2 µM FRAX597, black bars: 10 µM FRAX597. The graphs show means and standard deviations from 3 to 5 independent experiments for each cell line.

### Inhibitor toxicity

Toxicity of the inhibitors was determined after 5h treatment (Fig. 5A,B). Although increased numbers of apoptotic cells were observed after PIR3.5 treatment in some cell lines, they were always lower compared to samples treated with IPA-3, and the toxic effects of IPA-3 can thus be ascribed mostly to PAK inhibition. In the cell lines, FRAX597 usually induced less cell death than IPA-3 at the 5h time point (Fig. 5A), but the dead cell fraction increased after 22h (Fig. 5C). On the other hand, primary cells were much more sensitive to FRAX597 than to IPA-3 (Fig. 5B). No cytotoxic effects were detected after cell treatment with dasatinib.

We have noted markedly increased fluorescence signal in the Pacific blue channel in samples treated with FRAX597. FRAX597 fluorescence allowed us to follow the amount of this inhibitor in individual cells using flow cytometry. The distribution was largely heterogeneous, with at least two distinct subpopulations in all cell lines (examples of histograms are shown in Fig. 6). Correlation of FRAX597 signal with that of propidium iodide showed that although the dead cell fraction contained more FRAX597-high cells compared to live cells, both FRAX597-low and FRAX597-high subpopulations were present among dead cells (Supplementary Fig. S2B). Cell death was thus induced even in cells with low FRAX597 content. On the other hand, not all the cells with high FRAX597 content were dead after 22h (Fig. 5C, right). Cell lines with lower pPAK1 levels (KG-1, MOLM-7, Kasumi-1, see Fig. 1) were more resistant to FRAX597 treatment (Fig. 5C).

**Figure 6:**
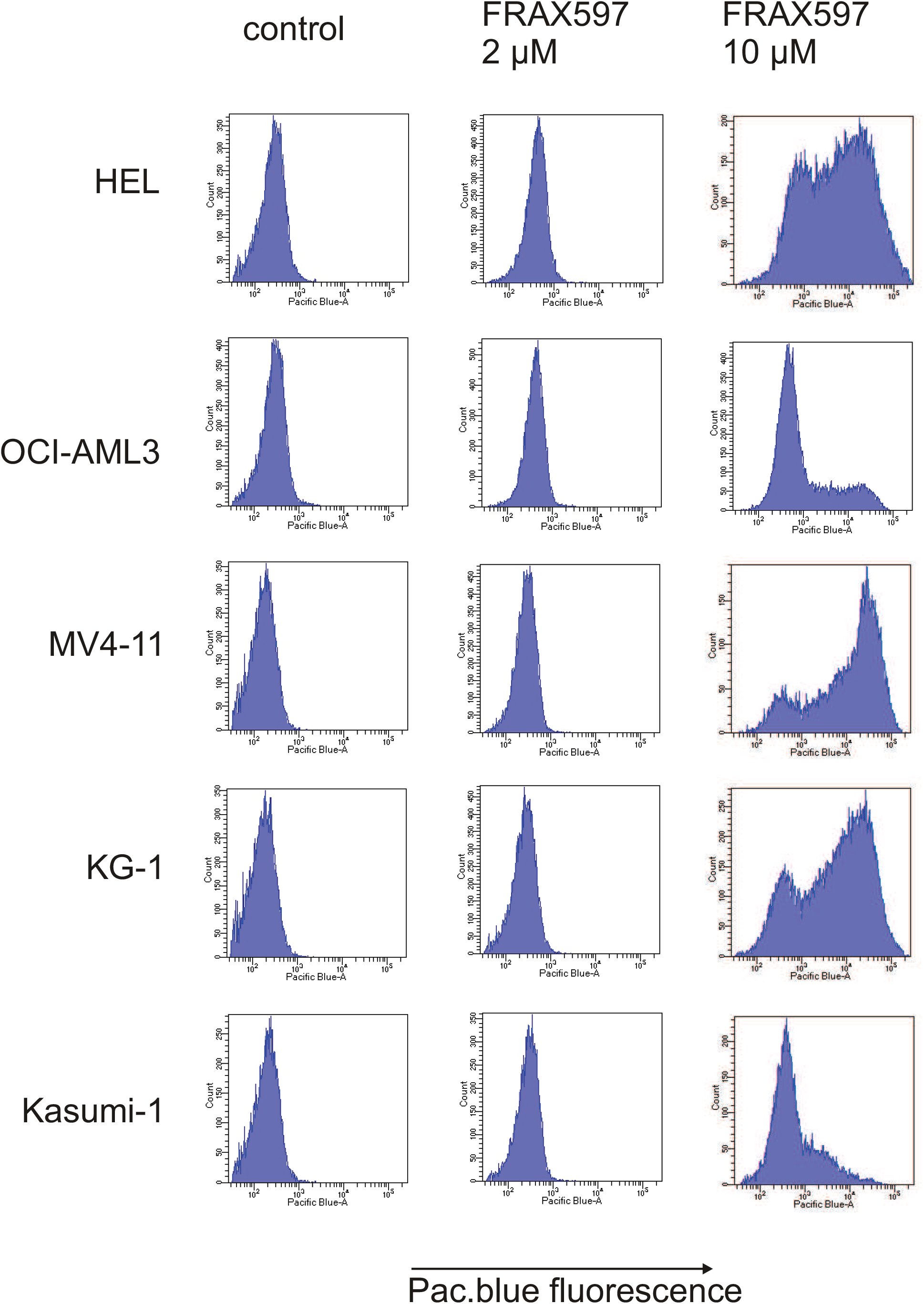
Examples of FRAX597 distribution in leukemia cells. Cells were treated with 2 or 10 µM FRAX597 and the fluorescence in the Pacific blue channel was measured after 5h incubation. Similar heterogenous distribution was observed in primary cells, as well as after 22h of treatment (see Supplementary Figure S2).

### Monitoring of cell-surface interaction by impedance measurement (ECIS)

To study the impact of PAK inhibition on cell-surface adhesion, we compared three cell lines with specific properties. OCI-AML3 cells were selected to represent AML blasts. Among AML cell lines, they have a high adhesivity to FN (about 80 %) and a high PAK1 and pPAK1 content (Figure 1C). MOLM-7 cells have high PAK2 levels (both transcript and protein, Figs 2 and 1C) and low-to-undetectable pPAK1. HEL cells express similar amount of PAK as OCI-AML3 (Fig. 1C), but they are known to be semi-adherent (37, 38) and differ from other leukemia cells in that they form more mature adhesion structures, which are regulated by Src kinase activity and by cytoskeletal tension (17).

Cell interaction with fibronectin was monitored by a non-invasive real-time method based on impedance measurement, ECIS (Electric Cell-Substrate Impedance Sensing). Changes of capacitance at high frequencies mainly reflect changes in the surface coverage, i.e. indicate the part of the surface that is covered with the cell membranes. At lower frequencies, the signal also depends on the proximity of the cell membrane to the surface. This indicates how tightly the cells are attached. In our experiments, the signal of resistance at low frequencies usually roughly mirrored the signal of capacitance at high frequencies (Supplementary Figure S3). However, the signal-to-noise ratio was higher and the changes induced by the inhibitors were more clear in the resistance at low frequencies. The resistance at 2 kHz was thus used to monitor the changes in cell interaction with the coated surface, in two different settings: the cells were either seeded into wells and treated after about 1.5 h, or pretreated for 1 h with the inhibitors and then seeded into FN-coated wells. Fig. 7 shows representative records obtained for HEL (panel A) and OCI-AML3 cells (panel B) in these two settings. Similar results were obtained in repeated experiments (at least 3 for each cell line and condition).

**Figure 7:**
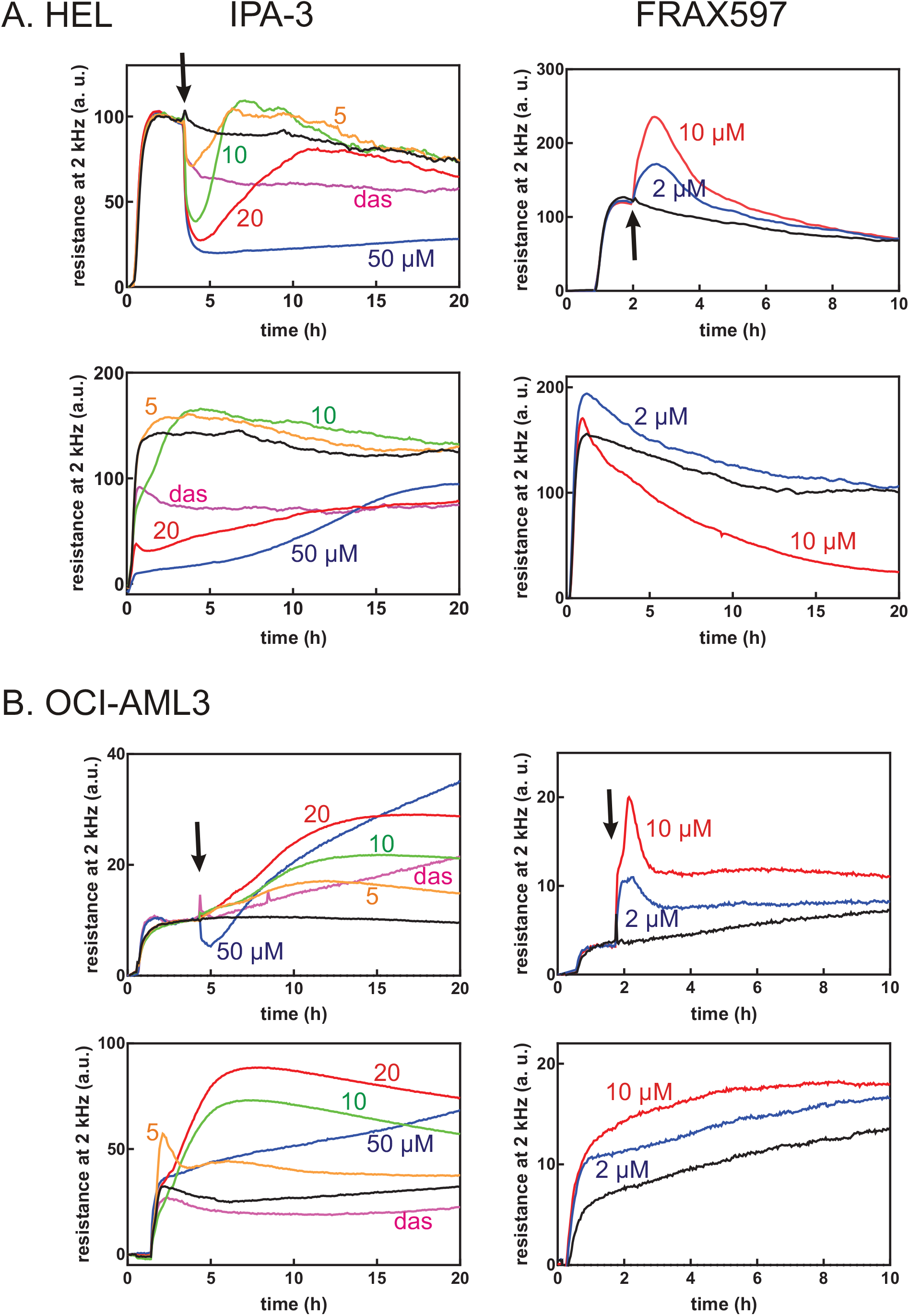
Effect of PAK inhibitors on cell interaction with fibronectin. Interaction of HEL (panel A) or OCI-AML3 (panel B) with fibronectin-coated surface was monitored through impedance measurement (ECIS). The cells were either allowed to adhere and subsequently treated with inhibitors (top images), or pretreated with inhibitors and seeded to coated wells (bottom images). The arrows mark the time of inhibitor addition. Untreated controls are represented by black lines. Left column: IPA-3 at 5-10-20-50 µM (orange-green-red-blue) and dasatinib 100 nM (magenta), right column: FRAX597 at 2-10 µM (blue-red). Representative records from at least 3 repeated experiments for each condition. The lines are means of two replicate wells, which were run in parallel.

In agreement with the adherent-like phenotype, HEL cell response to IPA-3 or FRAX597 was similar to that of adherent HEK-293T cells (29). In the first setting (Fig. 7A, upper row), a fast dose-dependent decrease of the resistance signal was induced by IPA-3, but not by FRAX597. Subsequently, a slower increase was observed at subtoxic IPA-3 doses, but not at 50 µM IPA-3. In contrast, FRAX597 addition resulted in a fast increase of the signal. In the pretreatment setting (Fig. 7A, bottom row), IPA-3 slowed down HEL cell attachment and reduced the amplitude of the signal, whereas FRAX597-pretreated cells attached as rapidly as the controls. On the other hand, no resistance drop was detected in OCI-AML3 cells after treatment with up to 20 µM IPA-3 and the signal progressively increased (Fig. 7B, top left). The effect of FRAX597 addition was similar to that in HEL cells (Fig. 7B, top right). As we have reported earlier, the fast drop upon IPA-3 treatment in adherent and semi-adherent cells was associated with cell contraction (29, 34). Microscopic examination of OCI-AML3 cells treated with FRAX597 confirmed that the increase in ECIS signal was due to an increase in the cell-surface contact area (Fig. 8). Closely similar results were obtained for MOLM-7 cells (Supplementary Fig. S4), which have low levels of PAK1/pPAK1.

**Figure 8:**
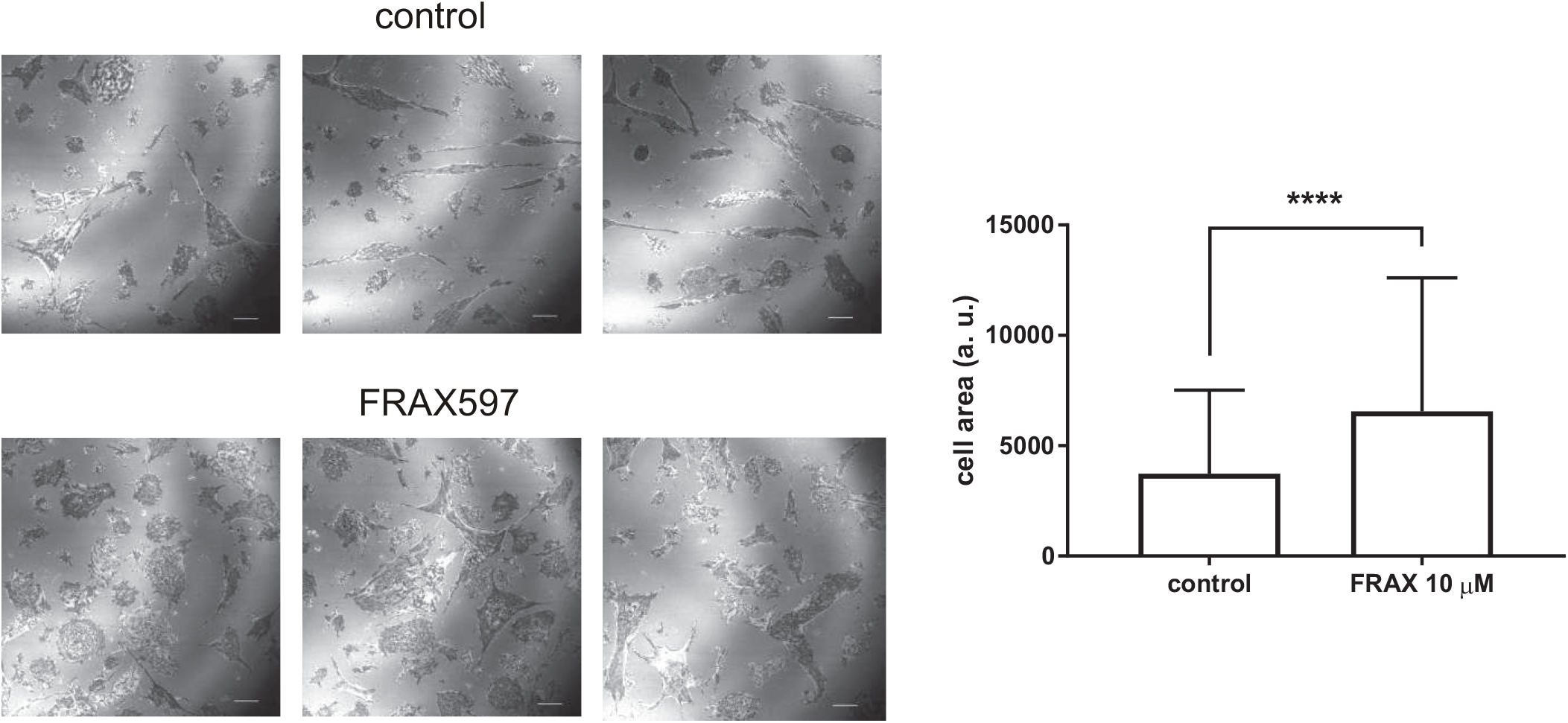
Microscopic analysis of cell-surface contact area in OCI-AML3 cells treated with FRAX597. OCI-AML3 cells were seeded on fibronectin-coated slide, incubated for 90 min and then treated for 30 min with 10 µM FRAX597. The cell parts closely adjacent to the surface were imaged using the interference reflection measurement. Representative slide sections are shown for control (top) and FRAX597-treated cells (bottom). Scale bar: 10 µm. Quantification of the cell-surface contact area from control cells (N = 184 cells from 3 biological replicates) and from cells treated with FRAX597 (N = 152 from 3 biological replicates) is shown on the right side, p < 0.0001 for the difference between the two groups by the standard two-tailed t-test.

The slow increase in the ECIS signal observed after IPA-3 addition to OCI-AML3 cells could be at least partly due to cell death induced by the treatment (Fig. 5A). Nevertheless, cell pretreatment with the pan-caspase inhibitor Q-VD-OPh (10 µM, added 30 min before IPA-3 treatment), which fully prevents caspase activation and apoptotic cell death, did not alter ECIS signal evolution (Supplementary Fig. S5).

### Adherent cell fraction measurement (ACF)

The stability of cell attachment to FN was evaluated by means of measurement of the adherent cell fraction (ACF) after 1h incubation on FN and subsequent wash. Fig. 9 shows the effect of 1h IPA-3 pretreatment for cell lines (panel A) and for primary cells (panel B). With regard to the heterogeneity of primary samples, we always assessed the phosphorylation status of Ser144/141 in these cells in parallel. Fig. 9B shows that ACF (solid lines) usually well correlated with the extent of PAK2 phosphorylation at Ser141 (dashed lines).

**Figure 9:**
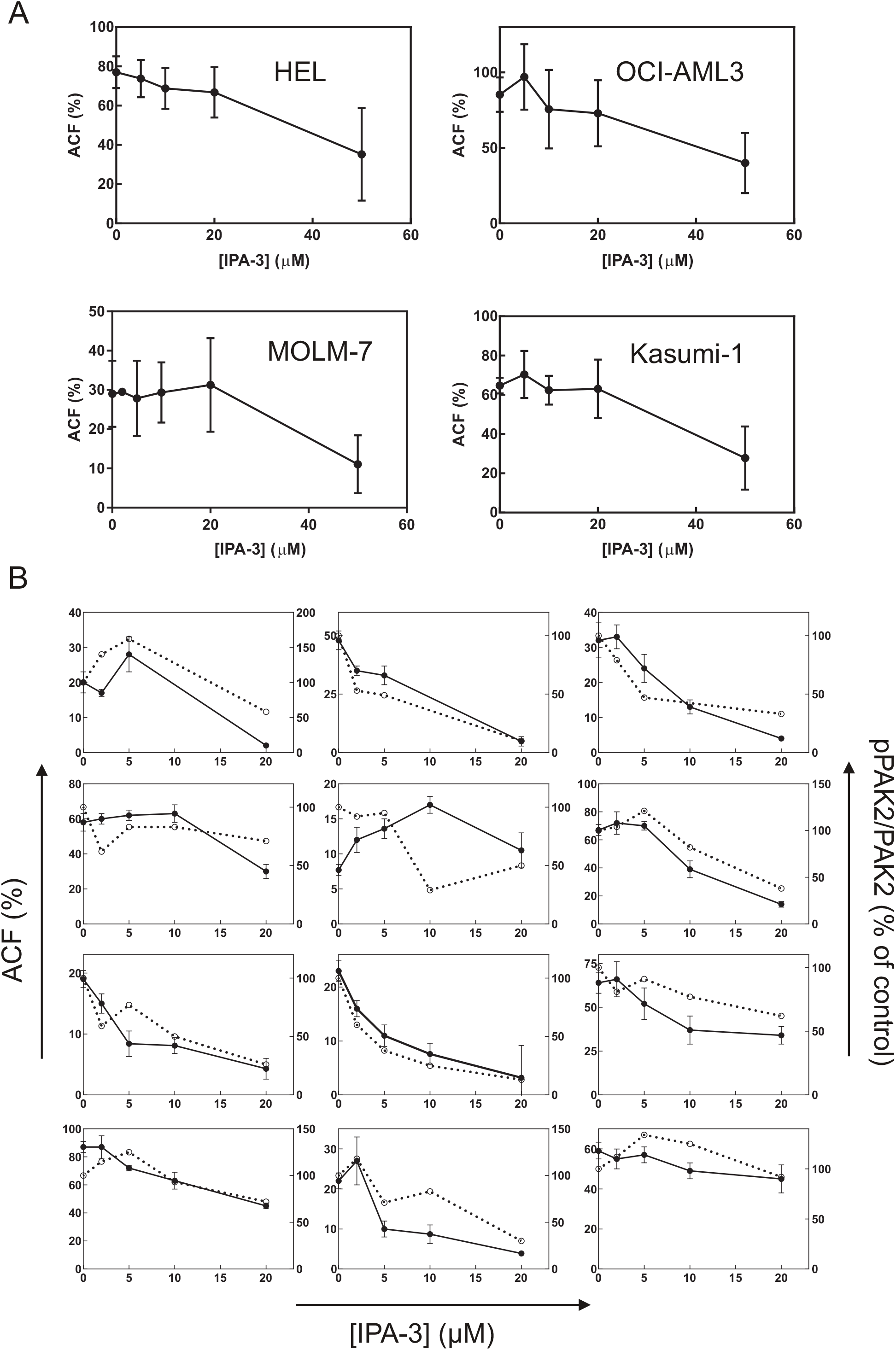
Effect of pretreatment with IPA-3 on the adhered cell fraction (ACF). Aliquots of cell lines (A) or of primary AML cells (B) were treated for 1h with IPA-3 at different concentrations as indicated and then seeded into fibronectin-coated wells. The fraction of attached cells was determined fluorometrically after 1h incubation. A: Means and standard deviations from 3 to 5 independent experiments are shown for each cell line. B: For primary AML cells isolated from 12 different patients, the extent of PAK2 phosphorylation at Ser144/141 was assessed by western-blotting in addition to ACF. The values of pPAK2 were normalized to total PAK2 and expressed as per cent of the controls (untreated). ACF: closed symbols and solid lines, left y-axis, pPAK2/PAK2: open symbols and dashed lines, right y-axis.

The effect of IPA-3 was then compared with that of PIR3.5, FRAX597 and dasatinib (Fig. 10). Except for MOLM-7 cells, IPA-3 treatment was associated with significant ACF drop whereas PIR3.5 had no or mild effect. Primary cells were somewhat more sensitive to all inhibitors compared to cell lines, i.e. lower concentrations were sufficient to induce comparable changes (Fig. 10B).

**Figure 10:**
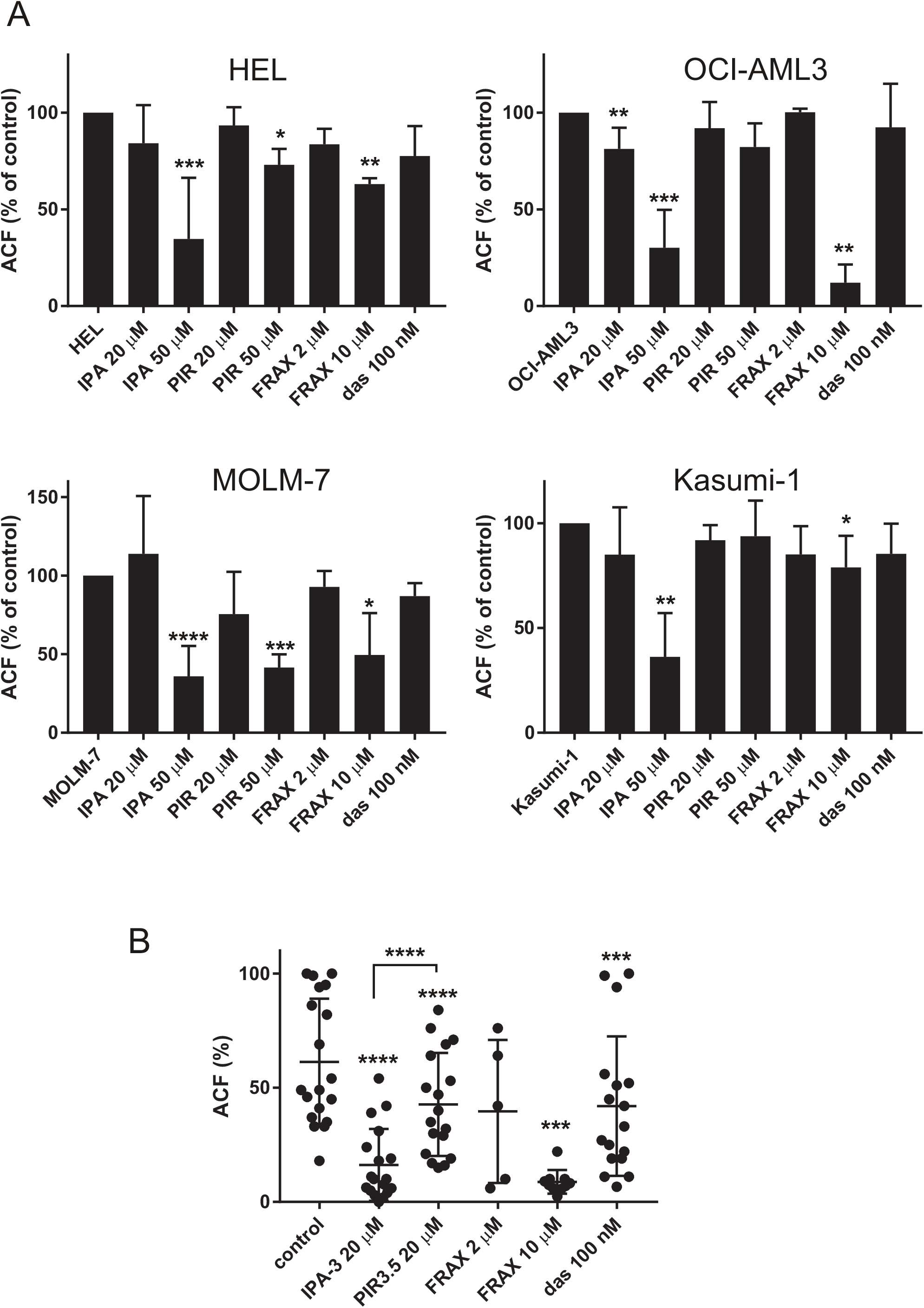
Effect of cell pretreatment with IPA-3, PIR3.5, FRAX597, or dasatinib on the adhered cell fraction (ACF). The cells were treated for 1h with the indicated inhibitor and then seeded to fibronectin. The fraction of attached cells was determined fluorometrically after 1h incubation. A: Summary results from 3 to 8 repeated experiments for each cell line, ACF is given as per cent of the corresponding controls. The bars show the means and standard deviations. B: Absolute ACF values from individual experiments with different primary AML cells. Differences between treated samples and the corresponding controls were evaluated using the standard paired t-test. * p less than 0.05, ** p less than 0.01, *** p less than 0.001, **** p less than 0.0001.

### FRAX597-induced changes in flow-cytometry scattergrams

Changes in cell morphology are often reflected in flow-cytometry scattergrams. We indeed noted that FRAX597 treatment altered flow cytometry scattergrams in all cell lines tested as well as in primary cells. An increase in side scatter (SSC) values was detected already after 1h cell treatment with 2 µM FRAX597 (Supplementary Fig. S6), and lasted for at least 24h.

### Correlation between PAK mRNA levels and CD34/integrin expression

Liu et al. reported different roles for PAK1-full and PAK1Δ15 in melanoma cell invasivity (39), which is usually associated with weaker cell-substrate adhesion. However, we found no clear correlation between PAK expression and AML cell adhesivity (Supplementary Fig. S7), although cells with very high adhesivity tend to have lower PAK1-full/PAK1Δ15 ratio. On the other hand, we noted a trend for negative correlation of this ratio with expression of CD34, a marker of stem and progenitor cells (Supplementary Fig. S7, bottom).

Cell adhesion to fibronectin is mainly mediated by the integrins α4β1, α5β1, and αVβ3, and we have recently described the expression of β1 and of αVβ3 integrins on primary AML cells in different AML subtypes (31). Correlation of integrin expression with PAK transcript levels showed statistically significant positive linear correlation of PAK1Δ15 and of PAK2, but not of the full-length PAK1 transcript, with both β1 and αVβ3 mean surface density (Supplementary Fig. S8).

## Discussion

PAK1 is a known mediator of c-Src, which promotes the epithelial-to-mesenchymal transition, adhesion structure dynamics, and cell movement. In cancer, Src activity is often associated with tumor invasivity, metastasis formation, and worse prognosis (40). PAK1 is involved in many different processes, including formation of membrane protrusions (41, 42), focal adhesion turnover (11-13), or regulation of cell contractility (43-46), although the individual functions were not attributed to specific PAK1 isoforms. Our recent results suggested that the full-length PAK1 mediates formation of membrane protrusions and enables spreading of adherent cells (29). On the other hand, the isoform PAK1Δ15 is to some extent similar to PAK2, and both PAK1Δ15 and PAK2 are found in focal adhesions and can thus play a role in formation, maturation, and/or disassembly of adhesion structures. Leukemia cells do not form mature focal adhesions, but the full-length PAK1 was present in membrane protrusions also in this cell type (Fig. 3). We have shown previously that leukemia cell binding to fibronectin-coated surface is not markedly affected by inhibition of Src kinases, and the Src/PAK1 axis is thus probably not important in this process. However, PAK2 was shown to be specifically required for homing of hematopoietic progenitors to the bone marrow (19). The importance of PAK2 in hematopoietic cells is further underlined by the fact that conditional PAK2 knock-out in hematopoietic stem cells induced multiple defects in cell differentiation affecting all the hematopoietic lineages (20).

Due to the complexity of western-blot PAK1 band pattern, it was not possible to discriminate between PAK1-full and PAK1Δ15 at the protein level (Fig. 1A). In addition, other transcription variants of PAK1 are annotated, and could also be translated. At the transcript level, we showed that the alternatively spliced variant(s) lacking the exon 15 (PAK1Δ15) is often more abundant than the full-length PAK1 (Fig. 2). Analysis of PAK transcripts also revealed differences between leukemia cell lines, which are established from leukemia stem cells, and bulk leukemia primary blasts, which are a mixture of many differentiation stages ranging from leukemia stem cells to more mature myeloid cells. Fig. 2 suggests that PAK1 variants, and especially the full-length PAK1, could be less expressed in leukemia stem cells. Accordingly, the ratio of PAK1-full to PAK1Δ15 transcripts inversely correlated with the fraction of primary cells expressing the stem and progenitor cell marker CD34 (Supplementary Fig. S7). On the other hand, PAK2 expression was more homogeneous and showed no difference between cell lines and primary cells. The prevalence of PAK2 in leukemia stem cells is in agreement with previous reports showing important roles for PAK2, but not for PAK1, in normal hematopoietic stem cells (19, 20).

Leukemia cells bind to the extracellular matrix, but the contact points are more dynamic and their composition corresponds to immature focal adhesions, which are not regulated by cytoskeletal tension (3, 17). Accordingly, adhesion of leukemia cell lines to fibronectin was not markedly affected by dasatinib, a potent Src kinase inhibitor ((17) ; Fig. 10). In contrast, dasatinib treatment lowered ACF in the majority of primary AML cells (Fig. 10B), which have higher PAK1 content (Fig. 2C). Although the stability of AML cell attachment to fibronectin did not depend on PAK levels (Supplementary Figure S7), we noted a positive correlation of PAK1Δ15/PAK2 transcript with the surface density of integrins (Supplementary Figure S8), which form the core of cell adhesion structures. This is in agreement with the localization of PAK1Δ15/PAK2 in adhesion structures of adherent cells (29), and indicates possible involvement of PAK1Δ15/PAK2 in the outside-in integrin signaling. Expression and Ser144 phosphorylation of un unspecified PAK1 isoform was indeed reported to be regulated by αVβ3 integrin in adherent cells (47).

In adherent cells, PAK1 inhibition by IPA-3, or indirectly by dasatinib, leads to rapid cell contraction, which can be detected through impedance monitoring (ECIS) (17, 29). A rapid resistance drop after IPA-3 or dasatinib addition was apparent also for the semi-adherent leukemia cell line HEL (Fig. 7A), but not for AML-derived OCI-AML3 cells (Fig. 7B), despite comparable amount of PAK (Fig. 1C). In contrast, FRAX597 induced a rapid increase in the resistance signal in both HEL and OCI-AML3 cell lines (Fig. 7, right column), in association with increased cell spreading (Fig. 8). In a previous report about adherent cells, the effects of IPA-3 and FRAX597 were similar: both inhibitors reduced cell stiffness, increased cell deformability and inhibited cell migration in 3D matrix (48). These consequences were probably mediated by changes in cell contractility and are thus not very relevant for leukemia cells. Moreover, in the above-mentioned work, the effects of inhibitors were tested after 2h incubation. Our experiments show that the immediate cell responses to IPA-3 and FRAX597 are rather opposite to each other. However, on a longer time scale, a slow increase in the resistance signal was present also upon IPA-3 treatment. We suggest that shortly after inhibitor addition, evolution of ECIS signal mainly reflects reduction of the kinase activity in the case of FRAX597 (36), and inhibition of non-kinase functions in the case of IPA-3 (35). Later on, a novel equilibrium is progressively achieved, and both inhibitors are thus expected to alter all PAK functions. Indeed, IPA-3 also induces PAK dephosphorylation at Ser144/141 after 1h treatment, in particular for the isoform denoted as pPAK1-low (Fig. 4). Interestingly, an *in vitro* PAK activation by Cdc42 was associated with a shift towards higher MW (11, 49), and the lower MW bands thus probably represent less active PAK forms, which are targeted by IPA-3. HEL cell contraction upon treatment with IPA-3 (Fig. 7A, top left) or with dasatinib is not mediated by inhibition of PAK kinase activity, as no signal drop was induced by FRAX597 (Fig. 7A). On the other hand, acute inhibition of PAK kinase activity is associated with an increase in the cell-surface contact area in both HEL and OCI-AML3 cells, and also in MOLM-7 cells (Supplementary Fig. S4), which have very low pPAK1 level (Fig. 1C). We thus speculate that the increased cell spreading is mainly due to inhibition of PAK2 kinase activity.

In the pretreatment setting (Fig. 7A and B, bottom graphs), IPA-3 slowed down cell attachment, supporting the view that PAK are required for formation of adhesion structures. Pretreatment with FRAX597 did not affect the attachment rate, and the kinase activity of PAK is thus likely dispensable for this process. The cells pretreated with FRAX597 even produced higher ECIS signal due to more extensive cell spreading (Fig. 8). However, despite higher ECIS signal, FRAX597 did not increase the adherent cell fraction (ACF, Fig. 10). At 2 µM concentration, FRAX597 slightly lowered ACF in some cell lines, but did not prevent cell binding, confirming that PAK kinase activity is not required for cell attachment to fibronectin. On the other hand, IPA-3 pretreatment was associated with dose-dependent reduction of cell adhesivity to fibronectin (Figs 9 and 10), despite more moderate effect on PAK phosphorylation in cell lines (Fig. 4). Higher FRAX597 doses also reduced the stability of cell attachment (Fig. 10), maybe due to a loss of non-kinase functions of PAK molecules with completely inhibited kinase domain. Alternatively, PAK kinase activity might be involved in adhesion structure stabilization. The dose-dependent changes of ACF nicely correlated with the extent of PAK2 phosphorylation in primary cells (Fig. 9B). These facts are in line with our previous finding that reduction of PAK2 expression by siRNA was associated with slower cell attachment in adherent cells (29).

In agreement with previous reports (5, 34), both IPA-3 and FRAX597 induced cell death in leukemia cells (Fig. 5), with different kinetics in the cell lines and primary cells. In particular, the toxic effect of FRAX597 was rapid in primary cells, but much slower in cell lines. This might be associated to higher PAK1 content in primary cells (Fig. 2), but also to inhibition of other target(s) as FRAX597 is less specific compared to IPA-3 (50). Interestingly, FRAX597-induced cell death was also fast in MV4-11 cells (Fig. 5A), which have an oncogenic mutation in the FLT3 kinase. The signaling from FLT3 was reported to be mediated by focal adhesion kinase (FAK) and PAK1 (23). FRAX597 distribution within cell populations was highly heterogenous (Fig. 6, Supplementary Fig. S2A), but dead cells were always present also in the subpopulation with low FRAX597 content (Supplementary Fig. S2B) and relatively low FRAX597 concentrations are thus sufficient to induce cell death. In cells treated with higher FRAX597 dose, the critical intracellular concentration was presumably achieved in a larger cell fraction, leading to higher percentage of dead cells in general. However, except for HEL cells, no increase in the dead cell fraction was observed at higher FRAX597 dose when only cells with high FRAX597 content were gated (Fig 5B, right). Thus, increasing the intracellular concentration beyond the critical limit does not enhance FRAX597 toxicity. Cells with low pPAK1 content (KG-1, MOLM-7, Kasumi-1) were less sensitive to FRAX597 (Fig. 5C), supporting the specificity of the observed effects. In line with our results, KG-1 cell line was previously shown to be more resistant to both IPA-3 and FRAX597 compared to three other AML cell lines (5).

Cell death induction could also contribute to the observed decrease in ACF, although the latter occurred quickly after inhibitor addition. PAK2, but not PAK1, is known to be cleaved during the apoptosis, the resulting fragment being constitutively active (51, 52). The cleaved PAK2 is required for extensive structural changes in cells undergoing apoptosis (53). In the case of IPA-3, a synergy of PAK inhibition with e.g. oxidative stress generation could contribute to larger inhibition of cell adhesion, especially in MOLM-7 cells and in primary AML cells, where the control compound PIR3.5 also lowered ACF (Fig. 10).

Requirement for PAK function in leukemia cells might also be responsible for the difficulties we experienced in leukemia cell transfection with plasmids for exogenous expression of PAK isoforms. Although we obtained reasonable yield of viable cells expressing the green fluorescent protein (GFP) from the control GFP-only plasmid, few cells survived after electroporation or nucleofection with GFP-labeled PAK. In adherent cells, the effect of PAK overexpression from these plasmids was similar to that of PAK silencing by siRNA, maybe due to inhibition of endogenous PAK by dimerization with the exogenous proteins (29). As the endogenous PAK are necessary for leukemia cell survival, their sequestration by the exogenous PAK could induce cell death.

We have also noted a rapid and durable increase in side scatter (SSC) values after cell treatment with FRAX597 (Supplementary Fig. S6). SSC is typically associated with cell granularity and, accordingly, FRAX597 was reported to induce formation of actin clusters in lamellipodia and of membrane-containing objects in the cytoplasm of adherent cells (48). However, the nature of these objects and the mechanism of their formation are unknown.

Fig. 11 sums up the current knowledge concerning the interplay between different members of PAK group I family, and illustrates our suggestions as to possible involvement of both kinase and non-kinase PAK activity in regulation of cell adhesion and survival. PAK1-full, PAK1Δ15, and PAK2 form homo- and heterodimers and are thus likely to mutually regulate their functions (29). Formation of membrane protrusions is promoted by kinase-independent function of PAK1-full (29), but it might be inhibited by PAK2 kinase activity, as FRAX597 increases the cell surface contact area in MOLM-7 cells with very low pPAK1 levels (Fig. S5). PAK non-kinase activity is required for adhesion structure assembly, since cell pretreatment with IPA-3, but not with 2 µM FRAX, prevents cell binding to fibronectin (Figs 9, 10). We have shown previously that both PAK2 and PAK1Δ15, but not PAK1-full, localize to adhesion structures in adherent cell types, and we thus suggest that PAK2 and/or PAK1Δ15 rather than PAK1-full might be involved in formation of cell-surface contact points. Finally, PAK inhibition induces cell death in leukemia cells. MOLM-7 with predominant PAK2 expression were more resistant to PAK inhibitors (Fig. 5) and PAK1 inhibition thus seem to dominate in this effect. In line with this hypothesis, reduction of PAK1, but not of PAK2 levels by means of shRNA was previously reported to inhibit cell proliferation in leukemia cell lines (5).

**Figure 11:**
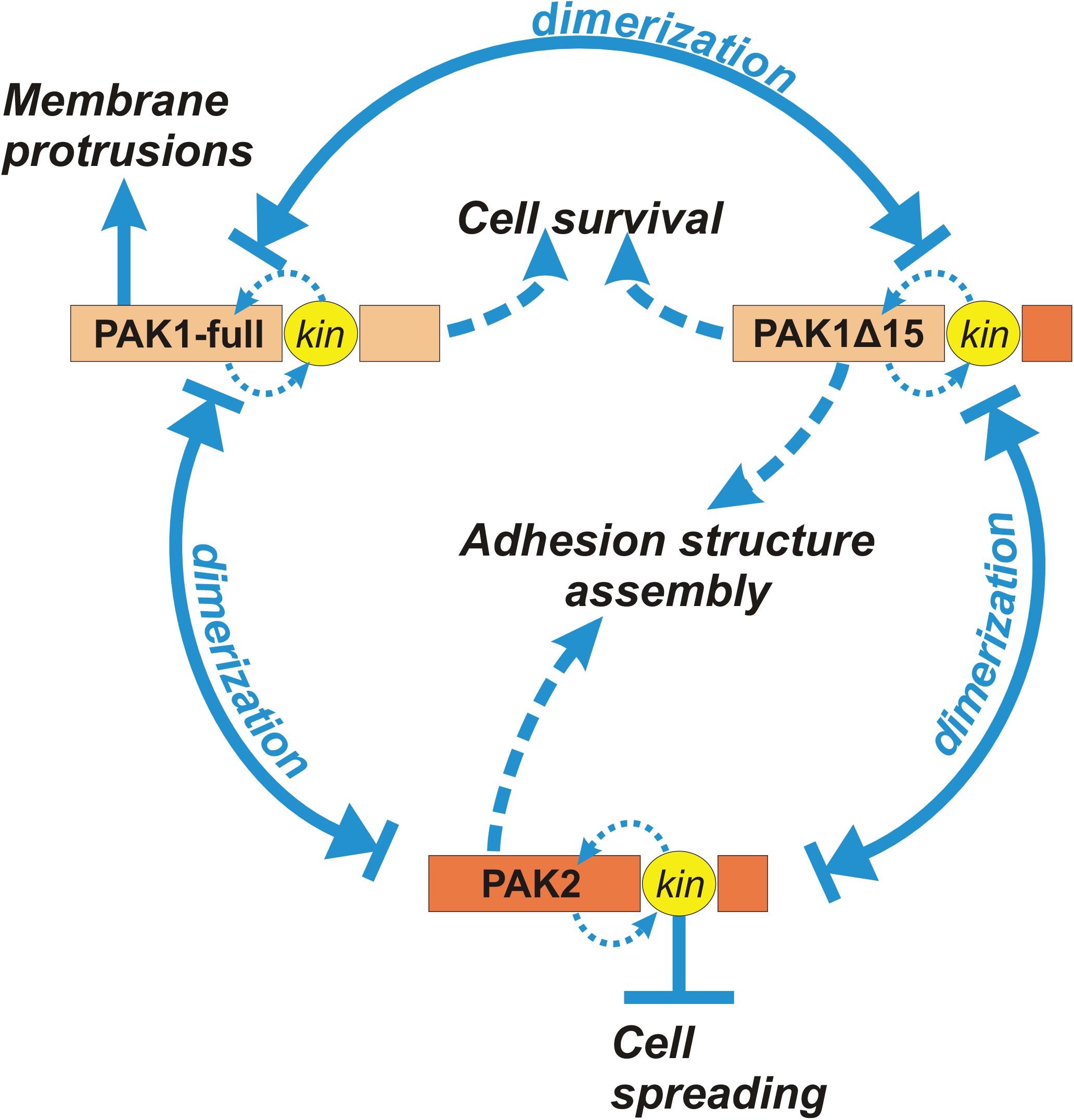
Scheme illustrating the complex interplay in PAK signaling. PAK1-full, PAK1Δ15, and PAK2 form homo- and heterodimers. Some of PAK functions are kinase-dependent, others are not. Blue arrows indicate reported or possible regulation of various cell processes by individual isoforms and/or domains (kinase versus non-kinase), suggested on the basis of our experimental results (see description in the text).

## Conclusions

Similarly to adherent cells, leukemia cells express both PAK1-full and PAK1Δ15. Both PAK1 isoforms are less abundant in leukemia cell lines compared to bulk primary leukemia cells, indicating that leukemia stem cells may express less PAK1 than more differentiated cells.

Although the adhesion structures of leukemia cells are different from those of adherent cells and are not so tightly regulated by kinases of the Src family, PAK are important for interaction of these cells with fibronectin. The inability to form adhesion points probably contributes to previously reported defects in engraftment of hematopoietic cells with PAK2 knock-out (19). The effects of acute inhibition of PAK kinase activity by FRAX597 were different from those induced by IPA-3, which preferentially targets kinase-inactive molecules. Inhibition of PAK kinase function in leukemia cells have similar consequences as increased Src activity in adherent cells, i.e. increased cell spreading and reduced adhesivity to the substrate. In addition, PAK inhibition by either IPA-3 or FRAX597 results in leukemia cell death.

Due to non-overlapping functions of PAK group I family members as well as of different PAK domains, possible therapeutic targeting of PAK will require attribution of individual functions to specific members and activities. In this context, knowledge about PAK family is still very limited and deserves attention. Tentatively, we propose a basic model, which is consistent with our experimental results as well as with previous reports (Fig. 11). However, many PAK functions remain to be associated with specific isoforms and type of activity.

## Acknowledgements

The authors wish to acknowledge M. Voráčová for technical assistance and Z. Gašová for providing cell samples from leukapheresis.

## References

1. Becker PS. Dependence of acute myeloid leukemia on adhesion within the bone marrow microenvironment. ScientificWorldJournal (2012) 2012:856467 doi: 10.1100/2012/856467; 10.1100/2012/856467.

2. Matsunaga T, Takemoto N, Sato T, Takimoto R, Tanaka I, Fujimi A, et al. Interaction between leukemic-cell VLA-4 and stromal fibronectin is a decisive factor for minimal residual disease of acute myelogenous leukemia. Nat Med (2003) 9:1158–65 doi: 10.1038/nm909.

3. Horton ER, Byron A, Askari JA, Ng DH, Millon-Fremillon A, Robertson J, et al. Definition of a consensus integrin adhesome and its dynamics during adhesion complex assembly and disassembly. Nat Cell Biol (2015) 17:1577–87 doi: 10.1038/ncb3257 [doi].

4. Ye DZ, Field J. PAK signaling in cancer. Cell Logist (2012) 2:105–16 doi: 10.4161/cl.21882 [doi].

5. Pandolfi A, Stanley RF, Yu Y, Bartholdy B, Pendurti G, Gritsman K, et al. PAK1 is a therapeutic target in acute myeloid leukemia and myelodysplastic syndrome. Blood (2015) 126:1118–27 doi: 10.1182/blood-2014-12-618801 [doi].

6. Semenova G, Chernoff J. Targeting PAK1. Biochem Soc Trans (2017) 45:79–88 doi: 10.1042/BST20160134 [doi].

7. Maruta H, Ahn MR. From bench (laboratory) to bed (hospital/home): How to explore effective natural and synthetic PAK1-blockers/longevity-promoters for cancer therapy. Eur J Med Chem (2017) 142:229–43 doi: S0223-5234(17)30565-2 [pii].

8. Zandvakili I, Lin Y, Morris JC, Zheng Y. Rho GTPases: Anti- or pro-neoplastic targets? Oncogene (2017) 36:3213–22 doi: 10.1038/onc.2016.473 [doi].

9. Kumar R, Sanawar R, Li X, Li F. Structure, biochemistry, and biology of PAK kinases. Gene (2017) 605:20–31 doi: S0378-1119(16)30981-7 [pii].

10. Rane CK, Minden A. P21 activated kinase signaling in cancer. Semin Cancer Biol (2018) doi: S1044-579X(17)30249-3 [pii].

11. Manser E, Huang HY, Loo TH, Chen XQ, Dong JM, Leung T, et al. Expression of constitutively active alpha-PAK reveals effects of the kinase on actin and focal complexes. Mol Cell Biol (1997) 17:1129–43.

12. Frost JA, Khokhlatchev A, Stippec S, White MA, Cobb MH. Differential effects of PAK1-activating mutations reveal activity-dependent and -independent effects on cytoskeletal regulation. J Biol Chem (1998) 273:28191–8.

13. Nayal A, Webb DJ, Brown CM, Schaefer EM, Vicente-Manzanares M, Horwitz AR. Paxillin phosphorylation at Ser273 localizes a GIT1-PIX-PAK complex and regulates adhesion and protrusion dynamics. J Cell Biol (2006) 173:587–9 doi: jcb.200509075 [pii].

14. Mayhew MW, Jeffery ED, Sherman NE, Nelson K, Polefrone JM, Pratt SJ, et al. Identification of phosphorylation sites in betaPIX and PAK1. J Cell Sci (2007) 120:o3911–8 doi: 120/22/3911 [pii].

15. Peri S, Navarro JD, Amanchy R, Kristiansen TZ, Jonnalagadda CK, Surendranath V, et al. Development of human protein reference database as an initial platform for approaching systems biology in humans. Genome Res (2003) 13:2363–71 doi: 10.1101/gr.1680803 [doi].

16. Renkema GH, Pulkkinen K, Saksela K. Cdc42/Rac1-mediated activation primes PAK2 for superactivation by tyrosine phosphorylation. Mol Cell Biol (2002) 22:6719–25 doi: 10.1128/mcb.22.19.6719-6725.2002 [doi].

17. Roselova P, Obr A, Holoubek A, Grebenova D, Kuzelova K. Adhesion structures in leukemia cells and their regulation by Src family kinases. Cell Adh Migr (2018) 12:286–98 doi: 10.1080/19336918.2017.1344796 [doi].

18. Kuželová K, Grebenová D, Holoubek A, Röselová P, Obr A. Group I PAK inhibitor IPA-3 induces cell death and affects cell adhesivity to fibronectin in human hematopoietic cells. PLoS ONE (2014) 9.

19. Dorrance AM, De Vita S, Radu M, Reddy PN, McGuinness MK, Harris CE, et al. The Rac GTPase effector p21-activated kinase is essential for hematopoietic stem/progenitor cell migration and engraftment. Blood (2013) 121:2474–82 doi: 10.1182/blood-2012-10-460709; 10.1182/blood-2012-10-460709.

20. Zeng Y, Broxmeyer HE, Staser K, Chitteti BR, Park SJ, Hahn S, et al. Pak2 regulates hematopoietic progenitor cell proliferation, survival and differentiation. Stem Cells (2015) doi: 10.1002/stem.1951 [doi].

21. Edlinger L, Berger-Becvar A, Menzl I, Hoermann G, Greiner G, Grundschober E, et al. Expansion of BCR/ABL1+ cells requires PAK2 but not PAK1. Br J Haematol (2017) doi: 10.1111/bjh.14833 [doi].

22. Phee H, Au-Yeung B, Pryshchep O, O’Hagan KL, Fairbairn SG, Radu M, et al. Pak2 is required for actin cytoskeleton remodeling, TCR signaling, and normal thymocyte development and maturation. eLife (2014) 3:e02270 doi: 10.7554/eLife.02270.

23. Chatterjee A, Ghosh J, Ramdas B, Mali RS, Martin H, Kobayashi M, et al. Regulation of Stat5 by FAK and PAK1 in Oncogenic FLT3- and KIT-Driven Leukemogenesis. Cell Rep (2014) 9:1333–48 doi: 10.1016/j.celrep.2014.10.039 [doi].

24. Berger A, Hoelbl-Kovacic A, Bourgeais J, Hoefling L, Warsch W, Grundschober E, et al. PAK-dependent STAT5 serine phosphorylation is required for BCR-ABL-induced leukemogenesis. Leukemia (2014) 28:629–41 doi: 10.1038/leu.2013.351 [doi].

25. Friedbichler K, Kerenyi MA, Kovacic B, Li G, Hoelbl A, Yahiaoui S, et al. Stat5a serine 725 and 779 phosphorylation is a prerequisite for hematopoietic transformation. Blood (2010) 116:1548–58 doi: 10.1182/blood-2009-12-258913 [doi].

26. Rennefahrt UE, Deacon SW, Parker SA, Devarajan K, Beeser A, Chernoff J, et al. Specificity profiling of Pak kinases allows identification of novel phosphorylation sites. J Biol Chem (2007) 282:15667–78 doi: M700253200 [pii].

27. Coniglio SJ, Zavarella S, Symons MH. Pak1 and Pak2 mediate tumor cell invasion through distinct signaling mechanisms.Mol Cell Biol (2008) 28:4162–72 doi: 10.1128/MCB.01532-07.

28. Lee JH, Wittki S, Brau T, Dreyer FS, Kratzel K, Dindorf J, et al. HIV Nef, paxillin, and Pak1/2 regulate activation and secretion of TACE/ADAM10 proteases. Mol Cell (2013) 49:668–79 doi: 10.1016/j.molcel.2012.12.004 [doi].

29. Grebenova D, Holoubek A, Roselova P, Obr A, Brodska B, Kuzelova K. PAK1, PAK1Delta15, and PAK2: similarities, differences and mutual interactions. Sci Rep (2019) 9:17171–6 doi: 10.1038/s41598-019-53665-6 [doi].

30. Tsuji-Takayama K, Kamiya T, Nakamura S, Matsuo Y, Adachi T, Tsubota T, et al. Establishment of multiple leukemia cell lines with diverse myeloid and/or megakaryoblastoid characteristics from a single Ph1 positive chronic myelogenous leukemia blood sample. Hum Cell (1994) 7:167–71.

31. Kuželová K, Obr A, Marková J, Gašová Z. Integrin expression and adhesivity to fibronectin in primary acute myeloid leukemia cells: Impact of NPM1 and FLT3 mutations. Eur J Haematol (2020) n/a doi: 10.1111/ejh.13488.

32. Kuzelova K, Pluskalova M, Brodska B, Otevrelova P, Elknerova K, Grebenova D, et al. Suberoylanilide hydroxamic acid (SAHA) at subtoxic concentrations increases the adhesivity of human leukemic cells to fibronectin. J Cell Biochem (2010) 109:184–95 doi: 10.1002/jcb.22397.

33. Kuzelova K, Grebenova D, Obr A, Roselova P, Holoubek A. Different roles for PAK1 and PAK2 in the interaction of leukemia cells with fibronectin. Febs J (2017) 284:354–5.

34. Kuzelova K, Grebenova D, Holoubek A, Roselova P, Obr A. Group I PAK inhibitor IPA-3 induces cell death and affects cell adhesivity to fibronectin in human hematopoietic cells. PLoS One (2014) 9:e92560 doi: 10.1371/journal.pone.0092560 [doi].

35. Deacon SW, Beeser A, Fukui JA, Rennefahrt UE, Myers C, Chernoff J, et al. An isoform-selective, small-molecule inhibitor targets the autoregulatory mechanism of p21-activated kinase. Chem Biol (2008) 15:322–31 doi: 10.1016/j.chembiol.2008.03.005.

36. Licciulli S, Maksimoska J, Zhou C, Troutman S, Kota S, Liu Q, et al. FRAX597, a small molecule inhibitor of the p21-activated kinases, inhibits tumorigenesis of neurofibromatosis type 2 (NF2)-associated Schwannomas. J Biol Chem (2013) 288:29105–14 doi: 10.1074/jbc.M113.510933 [doi].

37. Jarvinen M, Ylanne J, Vartio T, Virtanen I. Tumor promoter and fibronectin induce actin stress fibers and focal adhesion sites in spreading human erythroleukemia (HEL) cells. Eur J Cell Biol (1987) 44:238–46.

38. Ylanne J, Cheresh DA, Virtanen I. Localization of beta 1, beta 3, alpha 5, alpha v, and alpha IIb subunits of the integrin family in spreading human erythroleukemia cells. Blood (1990) 76:570–7.

39. Liu X, Si W, Liu X, He L, Ren J, Yang Z, et al. JMJD6 promotes melanoma carcinogenesis through regulation of the alternative splicing of PAK1, a key MAPK signaling component. Mol Cancer (2017) 16:175–2 doi: 10.1186/s12943-017-0744-2 [doi].

40. Guarino M. Src signaling in cancer invasion. J Cell Physiol (2010) 223:14–26 doi: 10.1002/jcp.22011.

41. Sells MA, Knaus UG, Bagrodia S, Ambrose DM, Bokoch GM, Chernoff J. Human p21-activated kinase (Pak1) regulates actin organization in mammalian cells. Curr Biol (1997) 7:202–10 doi: S0960-9822(97)70091-5 [pii].

42. Parrini MC, Camonis J, Matsuda M, de Gunzburg J. Dissecting activation of the PAK1 kinase at protrusions in living cells. J Biol Chem (2009) 284:24133–43 doi: 10.1074/jbc.M109.015271 [doi].

43. Chew TL, Masaracchia RA, Goeckeler ZM, Wysolmerski RB. Phosphorylation of non-muscle myosin II regulatory light chain by p21-activated kinase (gamma-PAK). J Muscle Res Cell Motil (1998) 19:839–54 doi: 10.1023/a:1005417926585 [doi].

44. Sanders LC, Matsumura F, Bokoch GM, de Lanerolle P. Inhibition of myosin light chain kinase by p21-activated kinase. Science (1999) 283:2083–5 doi: 10.1126/science.283.5410.2083 [doi].

45. Stockton RA, Schaefer E, Schwartz MA. P21-Activated Kinase Regulates Endothelial Permeability through Modulation of Contractility. J Biol Chem (2004) 279:46621–30 doi: 10.1074/jbc.M408877200.

46. Zeng Q, Lagunoff D, Masaracchia R, Goeckeler Z, Cote G, Wysolmerski R. Endothelial cell retraction is induced by PAK2 monophosphorylation of myosin II. J Cell Sci (2000) 113 (Pt 3):471–82.

47. Gasparski AN, Wilson JT, Banerjee A, Beningo KA. The Role of PAK1 in the Maturation of Invadopodia During Transient Mechanical Stimulation. Frontiers in Cell and Developmental Biology (2019) 7:269.

48. Mierke CT, Puder S, Aermes C, Fischer T, Kunschmann T. Effect of PAK Inhibition on Cell Mechanics Depends on Rac1. Front Cell Dev Biol (2020) 8:13 doi: 10.3389/fcell.2020.00013 [doi].

49. Manser E, Loo TH, Koh CG, Zhao ZS, Chen XQ, Tan L, et al. PAK kinases are directly coupled to the PIX family of nucleotide exchange factors. Mol Cell (1998) 1:183–92 doi: S1097-2765(00)80019-2 [pii].

50. Huynh N, Wang K, Yim M, Dumesny CJ, Sandrin MS, Baldwin GS, et al. Depletion of p21-activated kinase 1 up-regulates the immune system of APC(14/+) mice and inhibits intestinal tumorigenesis. BMC Cancer (2017) 17:431–0 doi: 10.1186/s12885-017-3432-0 [doi].

51. Lee N, MacDonald H, Reinhard C, Halenbeck R, Roulston A, Shi T, et al. Activation of hPAK65 by caspase cleavage induces some of the morphological and biochemical changes of apoptosis. Proc Natl Acad Sci U S A (1997) 94:13642–7.

52. Rudel T, Bokoch GM. Membrane and morphological changes in apoptotic cells regulated by caspase-mediated activation of PAK2. Science (1997) 276:1571–4.

53. Vilas GL, Corvi MM, Plummer GJ, Seime AM, Lambkin GR, Berthiaume LG. Posttranslational myristoylation of caspase-activated p21-activated protein kinase 2 (PAK2) potentiates late apoptotic events. Proc Natl Acad Sci U S A (2006) 103:6542–7 doi: 0600824103 [pii].

