## Supplementary Fig. for "Group I p21-activated kinases in leukemia cell adhesion to fibronectin"

Supplementary Figures

Figure S1: Western-blot examples, PAK dephosphorylation after 1h treatment with inhibitors

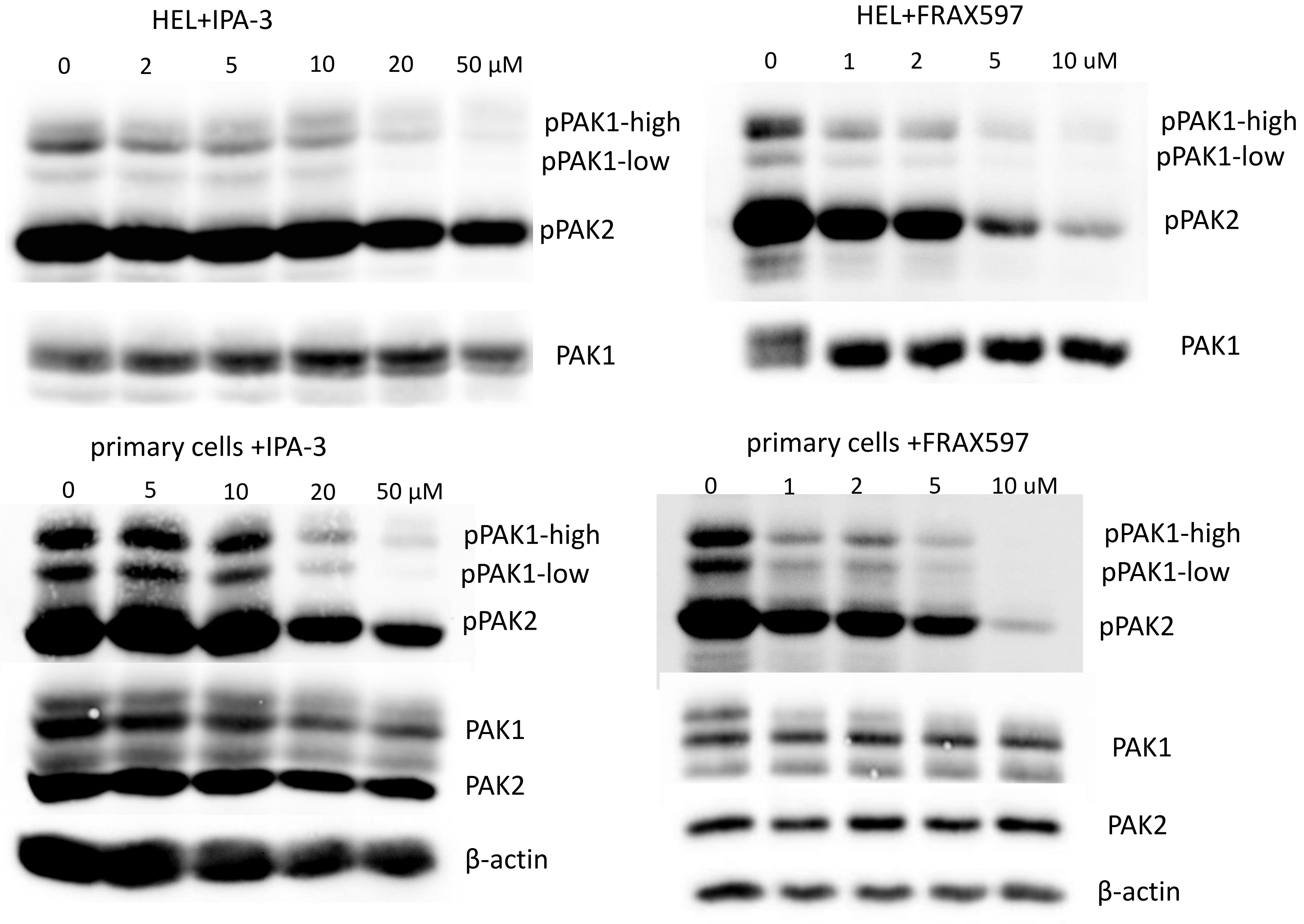

### Figure S2A: FRAX597 distribution in primary cells after 5h treatment

Mononuclear cell samples from three different AML patient were incubated for 5h with FRAX597 (2 or 10 µM) and the fluorescence intensity in the Pacific blue channel was measured using BD Fortessa flow cytometer.

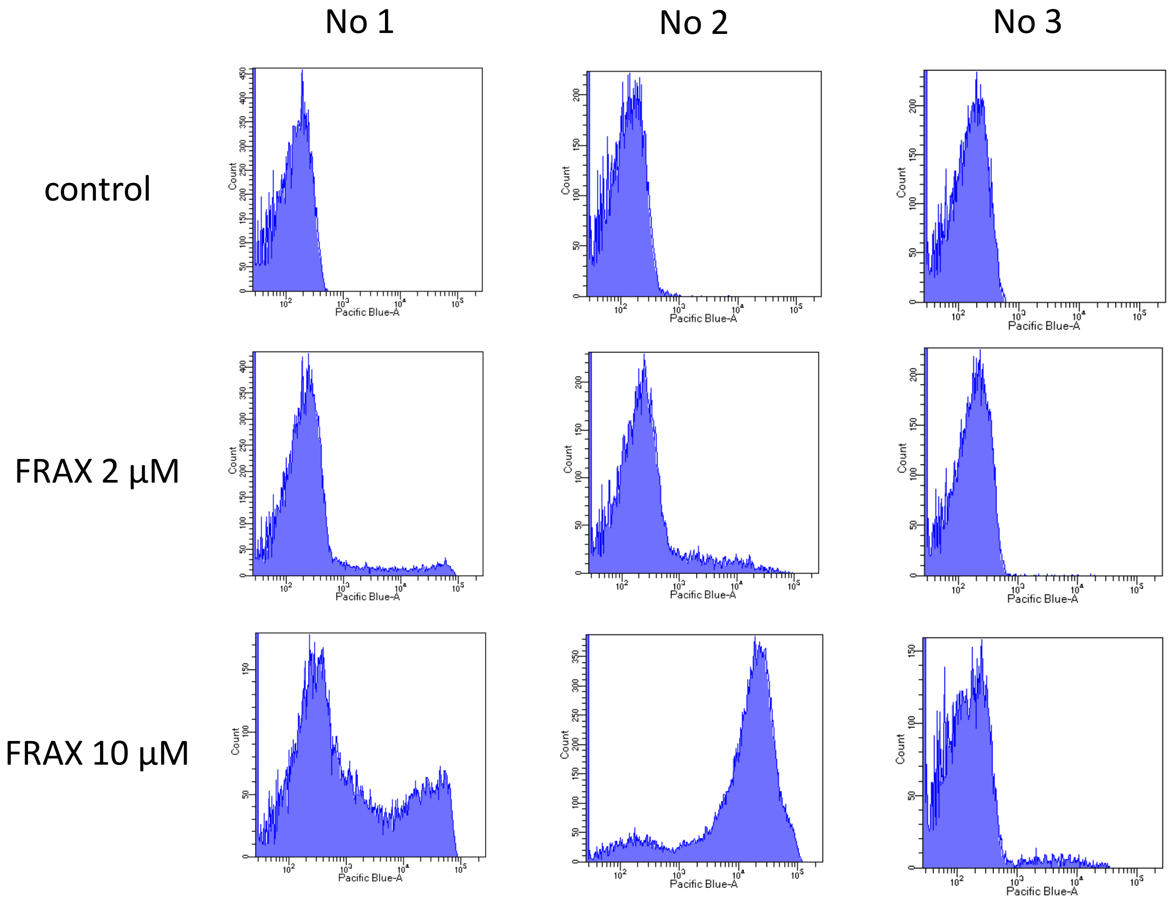

### Figure S2B: FRAX597 distribution in viable/dead cells after 22h treatment

Cells were incubated for 22h with FRAX597 (2 or 10 µM) and analyzed using BD Fortessa flow cytometer in the presence of propidium iodide (PI). The histograms show the fluorescence intensity in the Pacific blue channel for PI-negative cells (viable, blue) and PI-positive cells (dead, magenta).

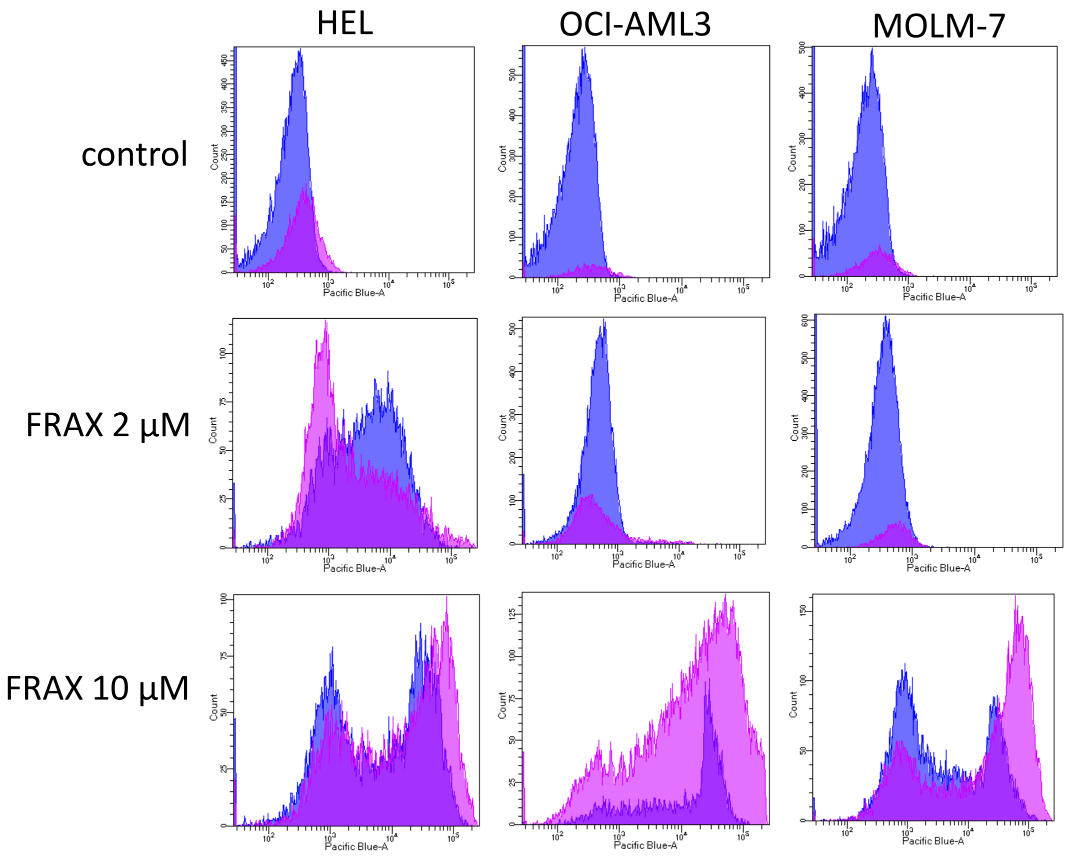

### Figure S3: Resistance and capacitance components of ECIS records

Comparison of ECIS signal course in the resistence at 2 kHz (top) and in the capacitance at 64 kHz (bottom).

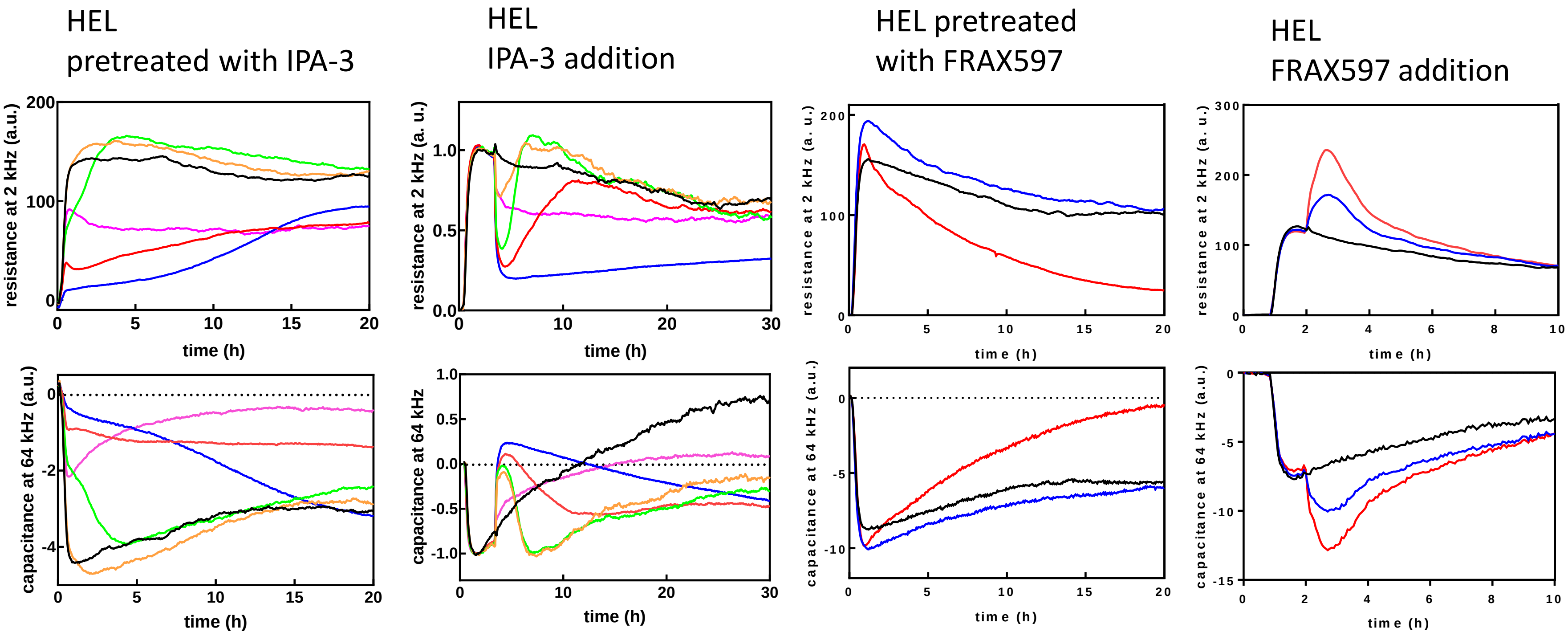

### Figure S3: Resistance and capacitance components of ECIS records - suite

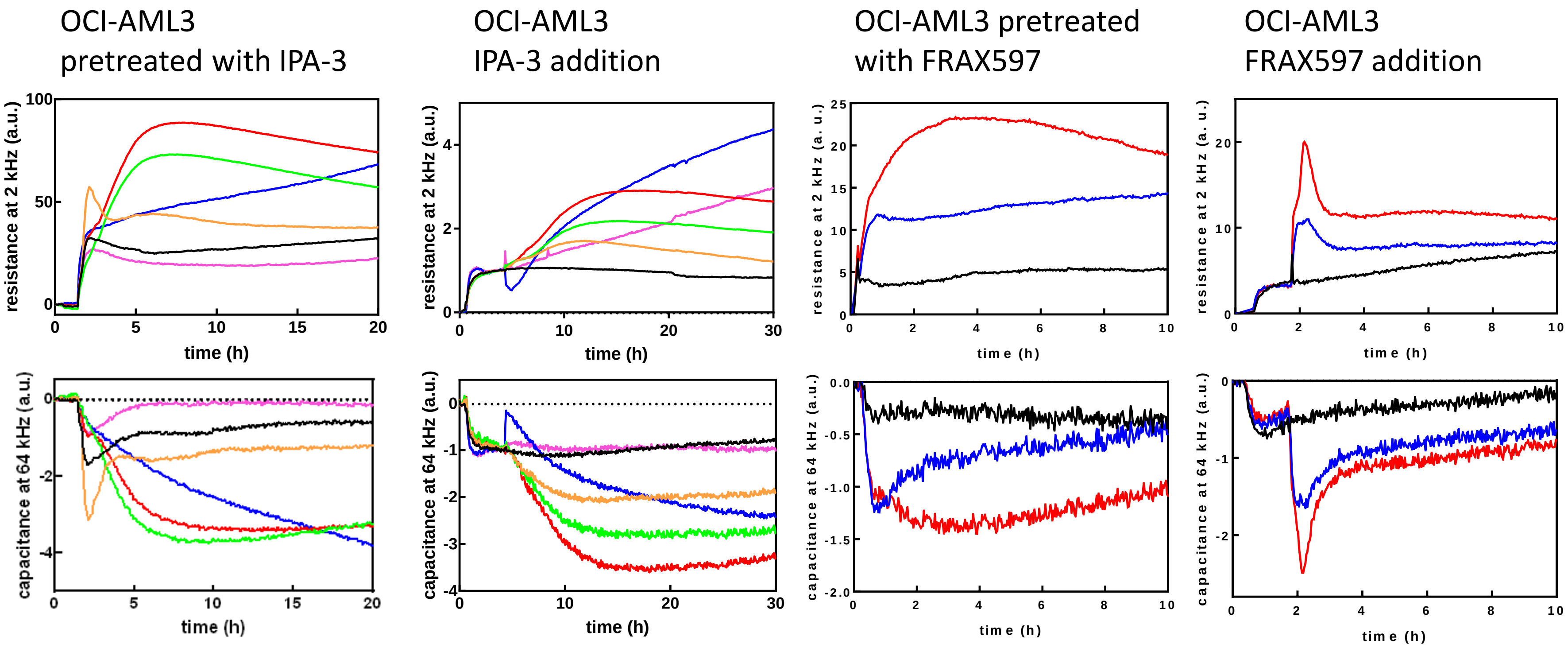

### Figure S4: Cell-contact area changes induced by FRAX597 in MOLM-7 cells

MOLM-7 cells were seeded on fibronectin-coated slides and treated for 30 min with 10 μM FRAX597. The contact area was visualized using the interference reflection measurement. Examples are shown on the left. Statistical evaluation of the cell area (from 116 control cells and 112 FRAX597-treated cells for randomly chosen views from 3 biological replicates) is shown on the right ($p<0.0001$).

control

FRAX597

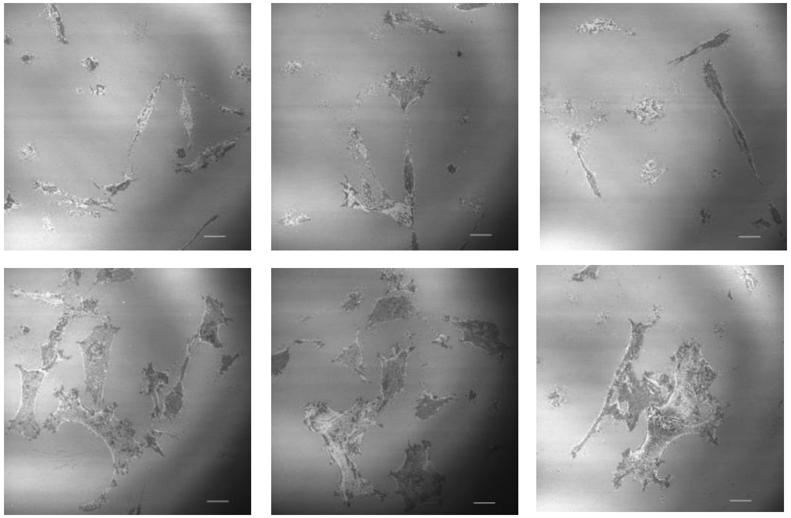

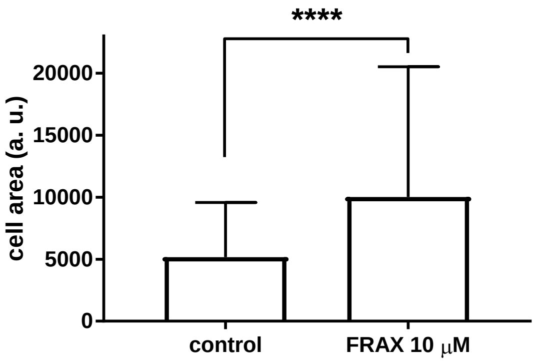


Representative ECIS records for MOLM-7 cells treated with FRAX597, 2 or 10 μM (blue-red).

Left: pretreatment for 1h
Right: FRAX597 addition (arrow)

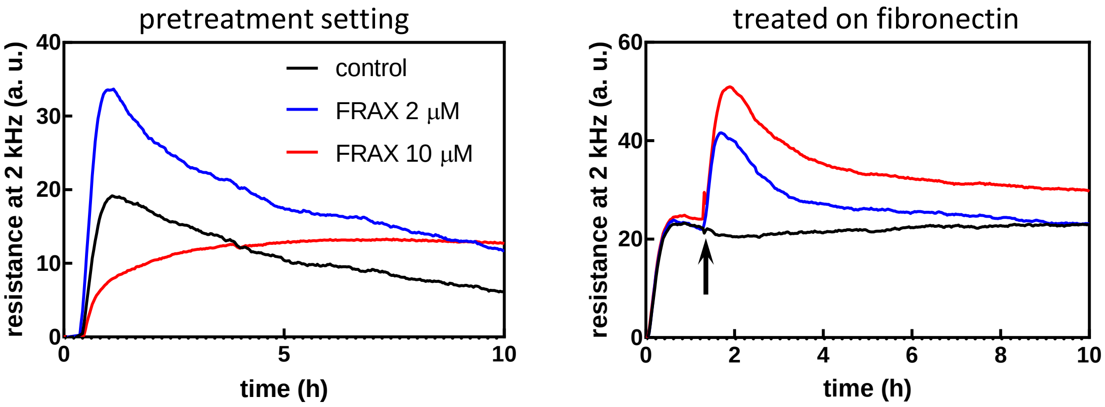

### Figure S5: Pretreatment with pan-caspase inhibitor does not affect ECIS signal

OCI-AML3 cells were seeded on fibronectin and monitored for about 1h before pretreatment with 10 µM Q-VD-OPh. IPA-3 (20 µM) was added after 30 min. The experiment was repeated with closely similar results.

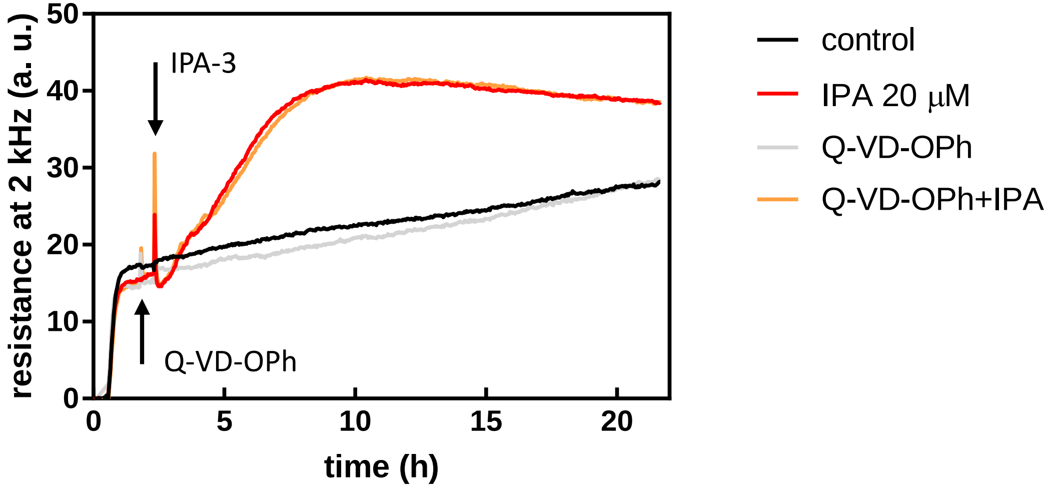

### Figure S6: FRAX597-induced changes in scattergrams

Top: Examples of scattergrams (forward scatter, FSC, versus side scatter, SSC) for OCI-AML3 after 1h treatment with 2 or 10 µM FRAX597. Cell debris and dead cells were outgated using FSC-low limit and PI-positivity, respectively. Bottom: Summary results for changes in the mean side-scatter (SSC) values after 1h treatment with FRAX597.

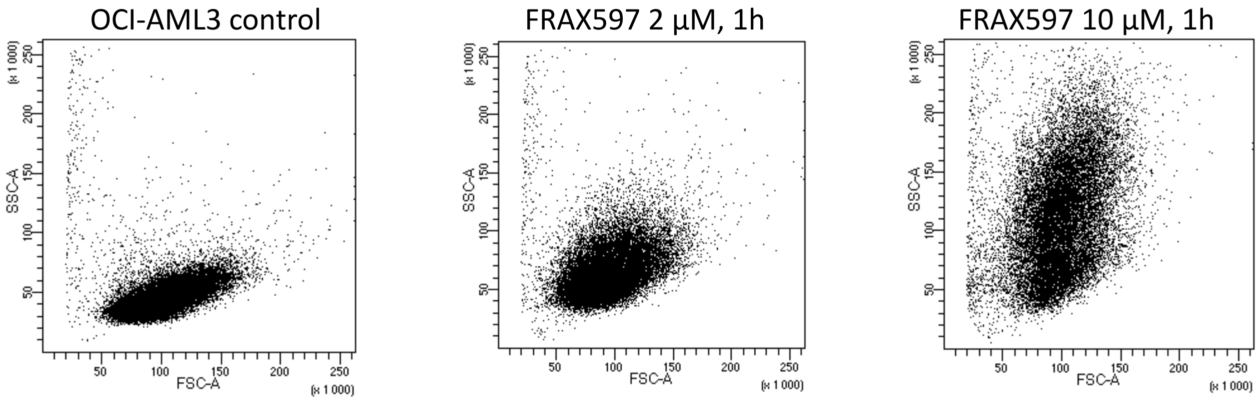


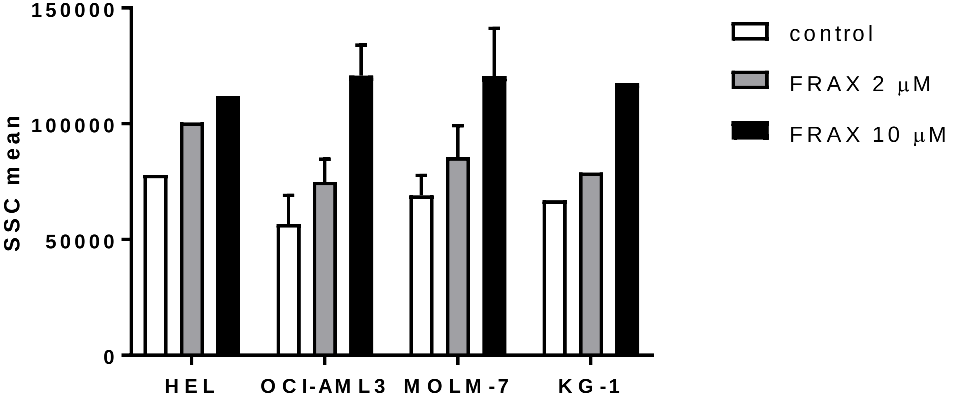

### Figure S7: PAK mRNA in primary AML cells correlated to cell adhesivity or to CD34 expression

Amounts of PAK1-full (green closed circles), PAK1Δ15 (red open circles) or PAK2 (blue squares) in primary AML cells were measured by RT-PCR, expressed as relative to GAPDH (left) or as ratios of PAK1 to PAK2 (middle) or PAK1-full to PAK1Δ15 (right), and plotted versus the adherent cell fraction (ACF, top) or versus the fraction of CD34-positive cells (determined by flow-cytometry from cryopreserved aliquots, bottom). CD34 is a marker of stem and progenitor hematopoietic cells. Pearson correlation coefficient and p-value from two-tailed t-test are given for the ratio PAK1-full/PAK1Δ15 mRNA.

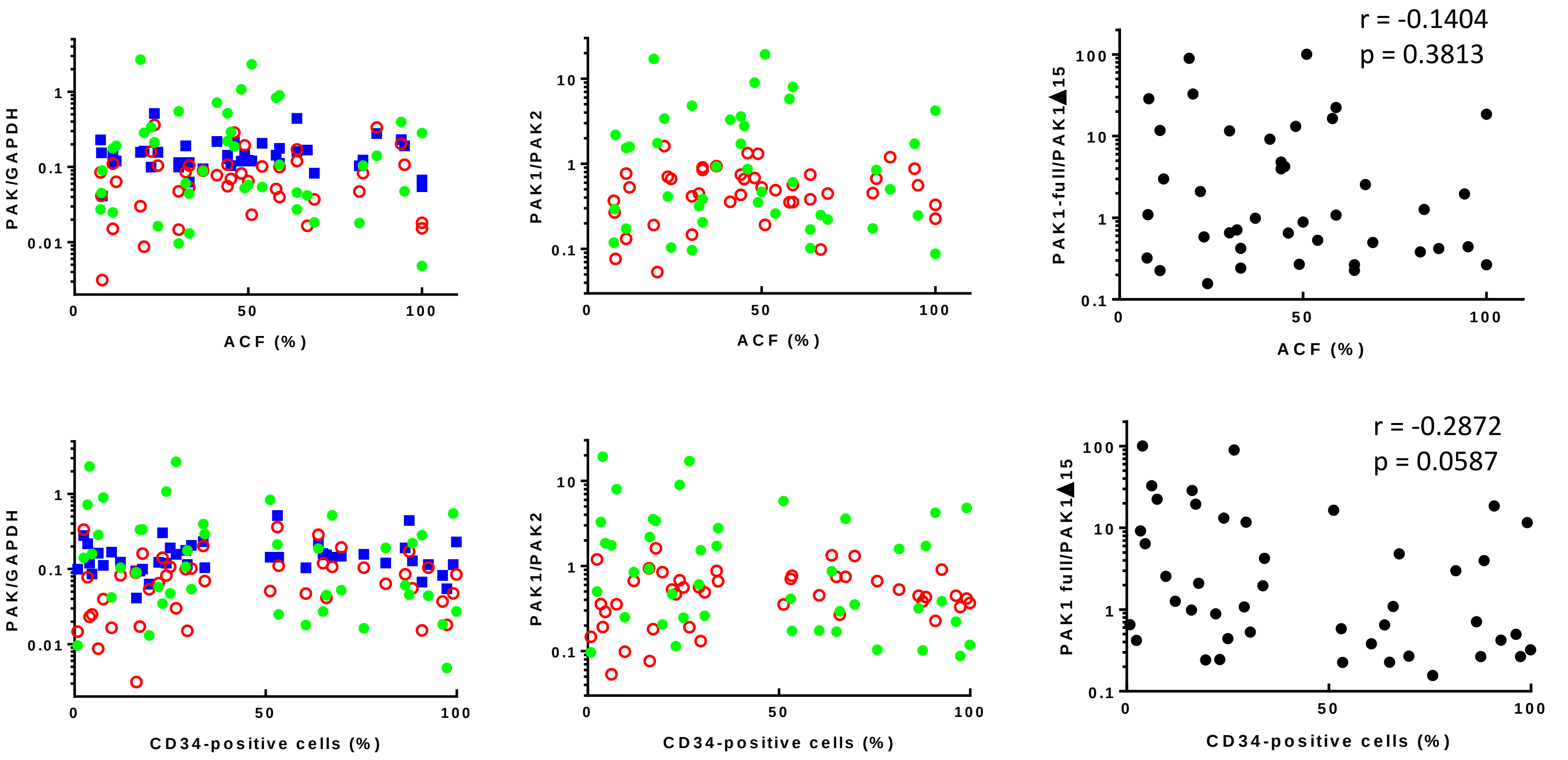

### Figure S8: PAK mRNA in primary AML cells correlated to integrin expression

Amounts of PAK1-full (green closed circles), PAK1Δ15 (red open circles) or PAK2 (blue squares) in primary AML cells were measured by RT-PCR, expressed as relative to GAPDH and plotted versus the mean surface density (MSD) of integrins β1 (left) or αVβ3 (right), which were determined by flow-cytometry (method described in Kuželová et al., European Journal of Haematology 2020, doi: 10.1111/ejh.13488). Correlation between transcript levels of PAK isoforms and integrin expression was assessed using GraphPad Prism 7 software (Pearson correlation test).

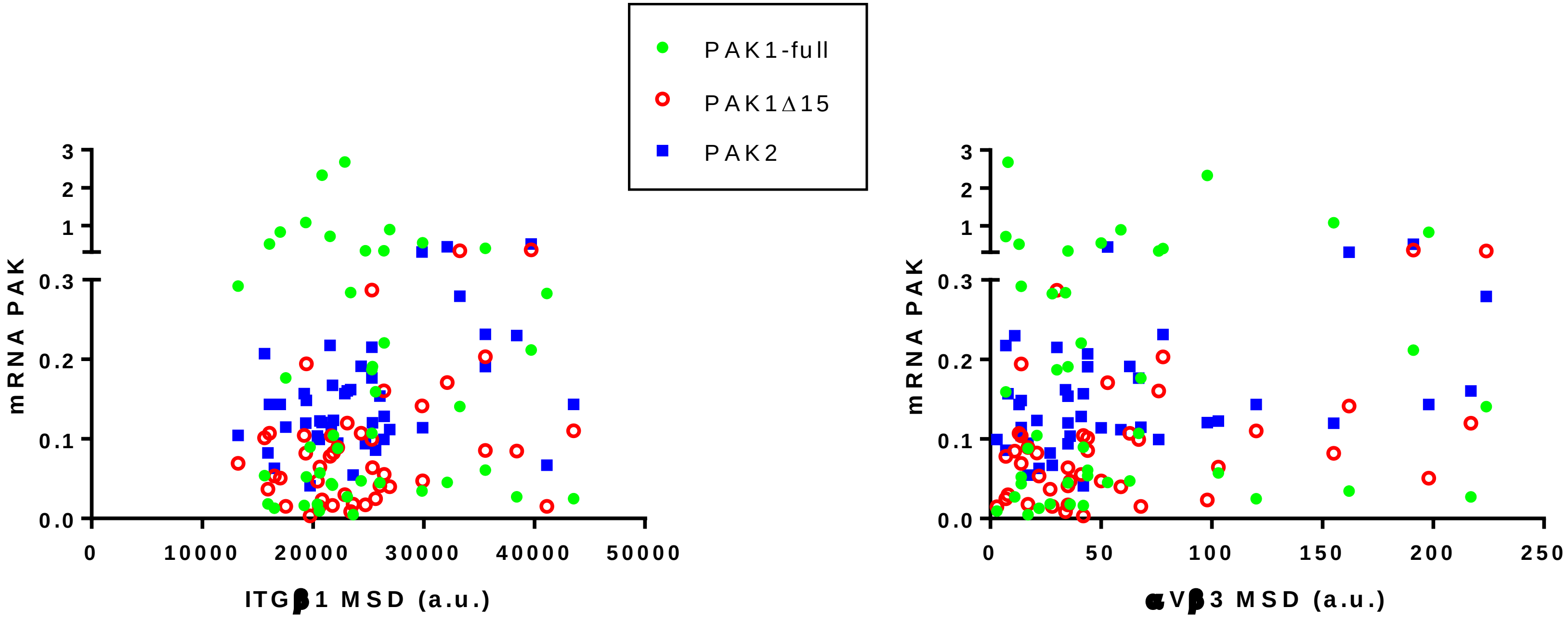


| PAK1-full length | PAK1Δ15 | PAK2 |
|---|---|---|
| r = -0.1163 | r = 0.3838 | r = 0.4641 |
| p = 0.4524 | *p = 0.0101 | **p = 0.0015 |

| PAK1-full length | PAK1Δ15 | PAK2 |
|---|---|---|
| r = 0.0568 | r = 0.4711 | r = 0.4224 |
| p = 0.7176 | **p = 0.0014 | **p = 0.0048 |
